# Translation in a Box: Orthogonal Evolution in the *Saccharomyces cerevisiae* Mitochondrion

**DOI:** 10.1101/2023.04.28.538752

**Authors:** Brooke Rothschild-Mancinelli, Claudia Alvarez-Carreño, Wenying Guo, Chieri Ito, Alex Costa, Anton S. Petrov, Kirill Lobachev, Loren Dean Williams

## Abstract

The ability to re-engineer and creatively evolve the translation system (TS) would allow invention of new coded polymers by altering the amino acid sidechain inventory and by shifting the polypeptide backbone into new chemical spaces. Unfortunately, the TS is difficult to manipulate and is more constrained over evolution than any other biological system. An orthogonal TS, running in parallel to the primary TS within a given host cell, would release constraints and allow TS manipulation. A fully orthogonal TS requires dedicated rRNAs, rProteins, aminoacyl-tRNA synthetases, and initiation and termination factors, none of which interact with the primary TS. The *S. cerevisiae* mitochondrial TS is fully orthogonal to the cytosolic TS. Mito-rRNAs, mito- rProteins, mito-tRNAs, mito-aminoacyl tRNA synthetases, and mito-translation factors are distinct from, physically separated from, and functionally independent of their cytosolic counterparts. Here, the *S. cerevisiae* mitochondrial translation system was subjected to various stresses including antibiotics, mutagenesis and truncation of mito-rProteins, or wholesale replacement of mito-rProteins. Directed evolution of these stressed systems was facilitated by controlled transitions between fermentation and respiration, by changing the carbon source in the growth medium; the dependence of *S. cerevisiae* survival on mitochondrial translation can be toggled on and off. Specific recreation of the resulting mutations recapitulate the evolved phenotypes. The method developed here appears to be a general approach for discovering functional dependencies. Suppressor mutations reveal functional dependencies within the *S. cerevisiae* mitochondrial TS. For example proteins Rrg9 or Mrx1 interact with the mito-TS and have critical role in its function. The combined results indicate that the *S. cerevisiae* mitochondrial TS can be engineered and evolved in isolation of the cytosolic TS.

**Significance:** The Central Dogma of Molecular Biology rules life on Earth. Information flows from DNA to mRNA to protein. In the last step of the Central Dogma, the translation system decodes mRNA and produces coded proteins by linking amino acids into polymers. Engineering and evolving the translation system could permits full technical control over this process and could lead to the generation of novel polymers. Here, we use the mitochondrial translation system in the budding yeast *Saccharomyces cerevisiae* for directed evolution of translation. We modify and evolve the translation system both directly and indirectly using antibiotics and gene editing tools and then measure resulting functionality. Our results show this secondary translation system inside *S. cerevisiae* mitochondria can be used as an approach for translation engineering.

## Introduction

The translation system (TS) is a central core of biology (1–3), reading mRNAs and writing out coded proteins. The ability to re-engineer and creatively evolve the TS would deepen our understanding of basic biology and lead to revolutionary advances in chemical and synthetic biology (4,5). Control over the TS would allow invention of new coded polymers by redrafting the protein sidechain inventory and by shifting the backbone into new chemical spaces (6,7). Production of encoded, functional, non-proteinaceous polymers would lead to revolutionary tools in catalysis, therapeutics, and materials.

Unfortunately, the TS is difficult to manipulate. The TS is more resistant to change over evolution than any other biological system (8–12). Around 90% of prokaryotic ribosomal RNA (rRNA) and 20 ribosomal proteins (rProteins) are universal in cytosolic ribosomes of all species. The three-dimensional structure of the core of the ribosome has been fixed for nearly 4 billion years (8). Even subtle changes to the TS are generally lethal. Further, with hundreds of proteins and thousands of nucleotides, the combinatorial mutation space of the TS is essentially insurmountable. Thus far, hese constraints have precluded technological control of the TS.

In principle, an orthogonal secondary TS, running in parallel to the primary TS within a given host cell, would release constraints and allow TS manipulation. The primary TS would keep house, producing the coded proteins required to maintain viability. Simultaneously, the secondary TS could be reengineered and subject to various evolutionary pressures (13,14). Changes in the secondary TS would not impact the primary TS or cell viability. TS orthogonality requires components that interact with each other but not with components of the primary TS (15). The demands for TS orthogonality appear immense. An orthogonal TS requires hundreds of dedicated macromolecules including rRNAs, tRNAs, rProteins, aminoacyl-tRNA synthetases, and initiation and termination factors that do not interact with those of the primary TS.

To our knowledge there are no previous descriptions of fully orthogonal TSs. Partial orthogonality has been established via distinct Shine-Dalgarno/anti-Shine-Dalgarno sequences, which inhibit interaction between primary mRNA/SSU RNAs and orthogonal mRNA/SSU RNAs (14). LSU rRNAs have been integrated into these orthogonal TSs by covalent linkage to orthogonal SSU rRNAs (16–18). In addition, some codons have been repurposed so that orthogonal aminoacyl-tRNA synthetase/cognate tRNA pairs direct incorporation of amino acids with non-canonical sidechains (7,19). In each of the partially orthogonal systems described above, rProteins, translation factors, most tRNAs and aminoacyl tRNA synthetases, and initiation and termination factors are shared between the primary and partially orthogonal TSs.

We are investigating a fully orthogonal TS contained within the mitochondrion of *Saccharomyces cerevisiae*. *S. cerevisiae* is a cheap, fast-growing and tractable organism with well-developed genetic tools. The *S. cerevisiae* mito-TS is fully orthogonal to the cytosolic TS. Mito-rRNAs, mito-tRNAs, mito-rProteins, mito-aminoacyl mito-tRNA amino-acyl synthetases, and mito-translation factors are distinct from, physically separated from, and functionally independent of their cytosolic counterparts. Further, the dependence of *S. cerevisiae* survival on mitochondrial translation can be toggled on and off. By changing the carbon source in the growth media, *S. cerevisiae* survive by either fermentation or respiration (20–22). Growth on glucose (fermentation) is independent of mitochondrial translation while growth on glycerol (respiration) requires mitochondrial translation. Once the mito-TS has been engineered or compromised, mutations can be accumulated under permissive conditions. A glucose medium (YPD) is permissive, and a glycerol medium (YPG) is non-permissive. Selection of suppressor mutants (those that suppress the non-functional phenotype) is achieved under non-permissive conditions.

Thus, genetic manipulation, treatment with small molecules and controlled transitions between fermentation and respiration allow directed evolution of the mito-TS. Here, the *S. cerevisiae* mito-TS was subjected to various stresses including antibiotics, mutagenesis and truncation of mito-rProtein, or wholesale replacement of mito-rProteins. Suppression of mito-translation disfunction was indicated by growth in non-permissive conditions and quantified by flow cytometry using a reporter strain in which mitochondrial translation produces mito-sfGFP (superfolder Green Fluorescence Protein) (23). We validated the results by recapitulating the evolved phenotype with specifically engineered suppressor mutations.

The combined results, including whole genome sequencing, indicate that the *S. cerevisiae* mitochondrial TS is orthogonal to the cytosolic TS and can be engineered and evolved. Suppressor mutations in uncharacterized proteins can reveal new information about dependencies within the *S. cerevisiae* mitochondrial TS.

## Results

Modified and compromised mito-TSs were optimized through accumulation and selection of compensatory mutations. Our approach can select for suppressors of TS disfunction in mito- rRNAs, mito-rProteins and in other auxiliary macromolecules associated with mito-translation. We monitored mito-TS function by ability to grow in restrictive media and by quantitating sfGFP production via flow cytometry in the MOY1355 strain (23). To our knowledge our approach to TS evolution has not been previously explored.

### Evolution of antibiotic resistance

As proof-of-concept, we used mito-TS evolution to restore function when *S. cerevisiae* mito-translation was inhibited by well-characterized small molecules. Our initial focus was on systems with predictable outcomes. We evolved resistance to mito-ribosome-targeting antibiotics chloramphenicol and erythromycin. We accumulated and selected mutations in the MOY1355 strain in the presence of chloramphenicol and erythromycin (Materials and Methods). We compared the suppressor mutations to previous mutations known to confer chloramphenicol and erythromycin resistance in bacterial and mito-ribosomes.

*Chloramphenicol*. Chloramphenicol binds to peptidyl transferase centers (PTC) and nascent polypeptide chains of bacterial and mito-ribosomes and prevents peptide bond formation (24,25). Chloramphenicol does not target cytosolic ribosomes of *S. cerevisiae* (26). Selective chloramphenicol inhibition of mito-ribosomes of the wild-type strain was confirmed here by lack of growth on non-permissive media (Figure 1a) and by absence of mito-sfGFP fluorescence, in the presence of chloramphenicol. The fluorescence intensity was equivalent to that of the non-fluorescent control (Figure 1c). Lack of inhibition of cytosolic ribosomes by chloramphenicol was confirmed by chloramphenicol-independent growth on permissive media (Supplementary Figure 1).

**Figure 1.**
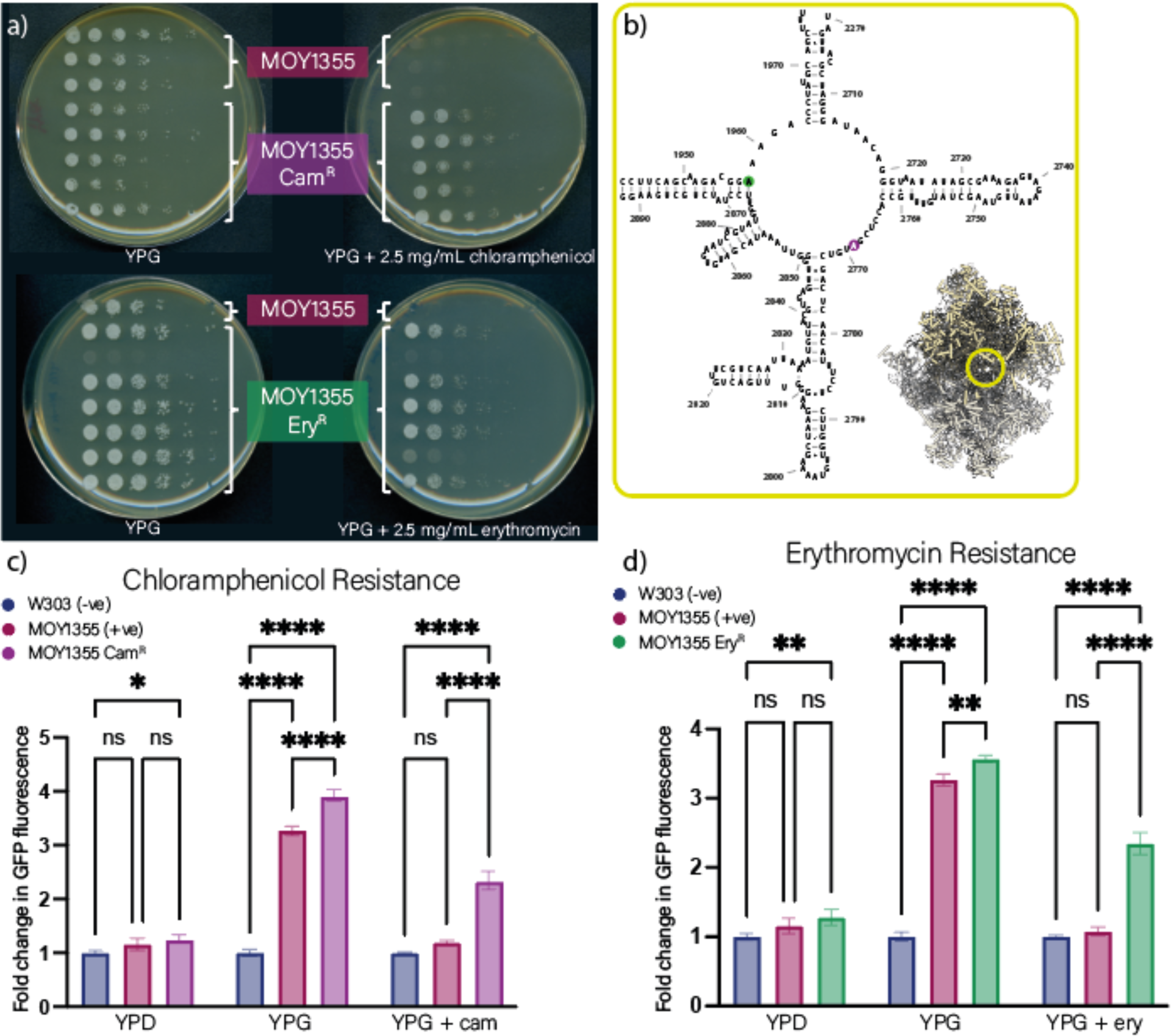
Mutations in mito-rRNA confer resistance to chloramphenicol and erythromycin. a) Growth of MOY1355, Cam^R^, and Ery^R^ on non-permissive (YPG), non-permissive + 2.5 mg/mL chloramphenicol, and non-permissive + 2.5 mg/mL erythromycin media. b) The location of the A2769C (A2503C) SNP in the 21S rRNA of the mito-ribosome in the Cam^R^ strain is indicated in the the 3D and 2D structure of the ribosome (PDB code 5MRC). c) and d) Function of the mito-translation system assayed by flow cytometry after growth in permissive (YPD), non-permissive (YPG), and non-permissive + antibiotics (YPG + 2.5 mg/mL chloramphenicol or erythromycin). W303 (blue) is the non-fluorescent negative control, MOY1355 (maroon) is the wild-type (WT) fluorescent positive control derived from W303 and the Cam^R^ strain (purple) Ery^R^ strain (green) are derived from MOY1355. Mean fluorescence intensity values were divided over W303 for the same experiment and media conditions to determine fold change over background (N = 10,000). ns = p > 0.05, * = p ≤ 0.05, ** = p ≤ 0.01, *** = p ≤ 0.001, *** = p ≤ 0.0001.

We evolved resistance to chloramphenicol (Cam^R^) in MOY1355. Suppressor mutations were accumulated during growth in permissive medium and selected by growth on non-permissive media containing chloramphenicol (Figure 1a). Sequencing revealed that Cam^R^ strains contain mutation A2769C (referred to as A2503C in *E. coli*), a SNP in the PTC of the LSU rRNA of the mito- ribosome (Supplementary Table 1; Figure 1b). The A2503C SNP has previously been shown to confer chloramphenicol resistance of mito-ribosomes and bacterial ribosomes (27). Fluorescence intensity indicates that mito-translation function of the Cam^R^ strain in non-permissive media is greater than that of wild-type MOY1355. Cell growth and mito-translational function of Cam^R^ mutants were maintained in non-permissive media in the presence of chloramphenicol at slightly reduced levels (Figure 1c).

#### Erythromycin

Erythromycin obstructs the exit tunnel of bacterial and mito-ribosomes but not cytosolic ribosomes of eukaryotes (24). As with chloramphenicol, erythromycin selectively inhibits mito-ribosomal function in *S. cerevisiae* (Supplementary Figure 2). We evolved resistance to erythromycin (Ery^R^) in MOY1355 via growth in permissive media to accumulate mutations, followed by selection for suppressors in non-permissive media containing erythromycin. Resistant mutants (Ery^R^) sustained erythromycin-independent growth in non-permissive media (Figure 1a). Wild-type MOY1355 grew in non-permissive media, only in the absence of erythromycin (Figure 1a). Sequencing of Ery^R^ isolates revealed an A1958G (referred to as A2508G in *E. coli*) SNP in the LSU mito-rRNA (Supplementary Table 1; Figure 1b). This A2508G SNP has been shown previously to confer resistance to erythromycin (28). Florescence intensity suggests that function of mito-translation of the Ery^R^ strain in non-permissive media is greater than that of wild-type MOY1355. Cell growth and mito-ribosomal function of Ery^R^ mutants were maintained in non-permissive media in the presence of erythromycin at slightly reduced levels. MOY1355 did not grow on non-permissive media in the presence of erythromycin; mito-translation function was similar to that of the non-fluorescent control (Figure 1d).

### Mito-rProtein Perturbations

The results with antibiotics (above) demonstrate that the mito-ribosome is an orthogonal TS that can be subject to directed evolution without perturbing the cytosolic (primary) TS. Thus, we investigated evolutionary adaptation to changes in mito-rProteins. The orthogonality of the cytosolic and mito-translation systems allows targeted perturbation of mito-rProteins without affecting cytosolic rProteins. We made various edits to sequences of mito-rProteins, which were incorporated into the mito-ribosome. These edits impaired mito-TS functionality. We evolved and sequenced suppressor strains with recovered mito-translational functionality.

One goal here was to manipulate one of the central and critical elements of the TS – the nascent polypeptide exit tunnel. Control over exit tunnel geometry may ultimately allow production of novel backbones (6). We used directed evolution to recover function of mito-ribosomes with perturbations of the exit tunnel. The canonical ribosomal exit tunnel is diverted in the *S. cerevisiae* mito-ribosomes (29,30) in part by an extension of mito-uL23 as well as by recruitment of mL50 (31).

#### Truncated mito-rProtein mL50

mL50_1-264_ (1-264 indicates the length of the native protein) is an integral part of the *S. cerevisiae* mito-exit tunnel (Figure 2a). mL50_1-264_ was truncated here by the removal of part of the C-terminal α-helix (aa 250-264) to create mL50_1-249_ within the MOY1355 strain. It was anticipated that mL50_1-249_ would significantly impair mito-translation. Indeed, the mL50_1-249_ strain showed decreased mito-ribosomal function compared to wild-type MOY1355. Mito-function decreased from 3.3-fold over background to 2.3-fold over background as suggested by mito-sfGFP fluorescence (Figure 2c).

**Figure 2.**
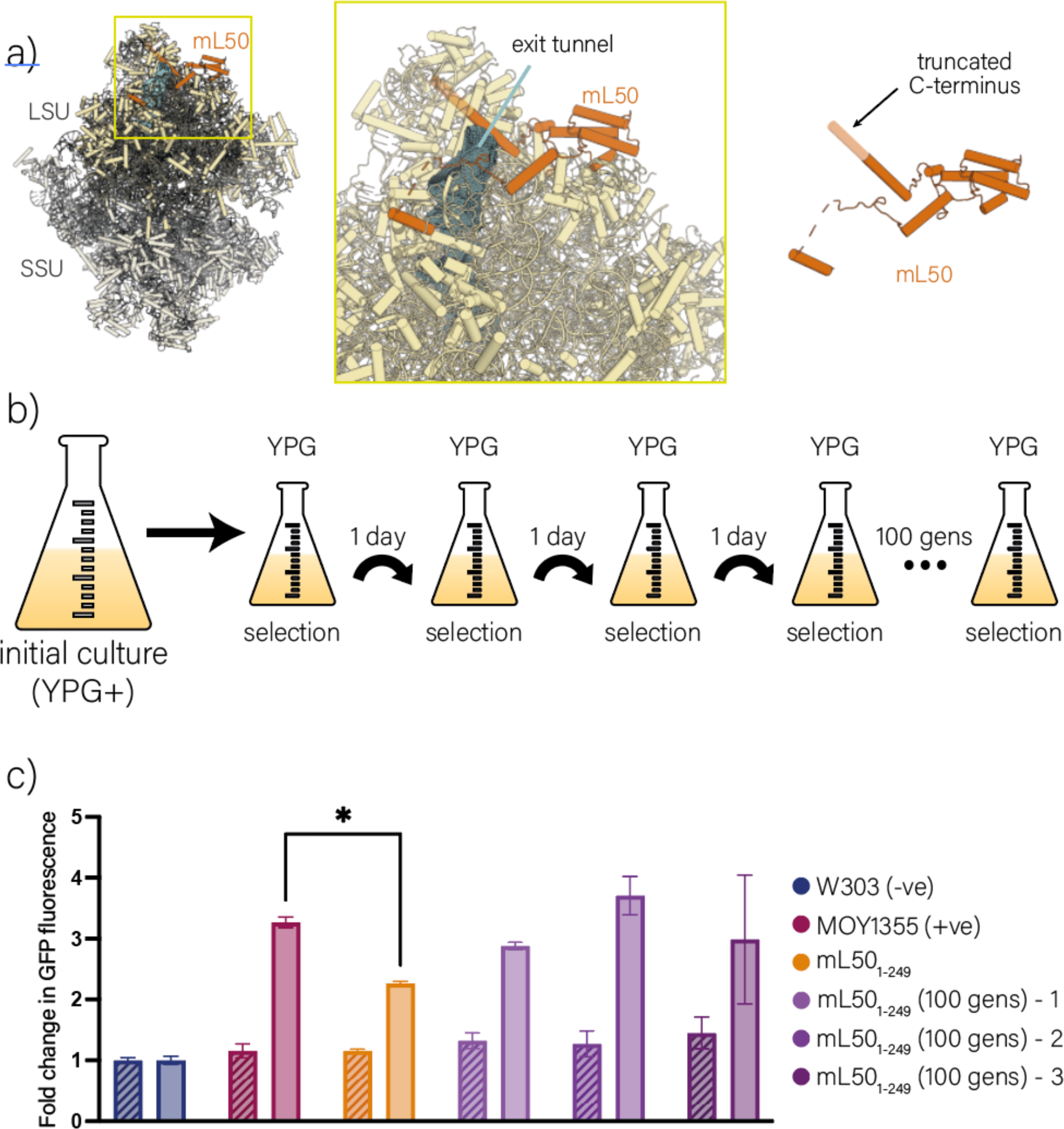
TS function can be recovered via ALE. a) mito-TS function was compromised by engineered truncation of the protein mL50 (orange) (PDB code 5MRC). The nascent peptide exit tunnel is indicated. b) ALE for 100 generations in non-permissive (YPG) media was used to recover mito-TS function. c) The function of the mito-TS was assayed via flow cytometer after growth in permissive (YPD; diagonal stripes) and non-permissive (YPG; solid pattern) media. W303 (blue) is the non-fluorescent negative control, MOY1355 (maroon) is the WT fluorescent positive control derived from W303 and mL50_1-249_ (orange) is the initial truncated strain with its evolved counterparts in shades of purple. Mean fluorescence intensity values were divided over W303 for the same experiment and media conditions to determine fold change over background (N = 10,000). ns = p > 0.05, * = p ≤ 0.05, ** = p ≤ 0.01, *** = p ≤ 0.001, *** = p ≤ 0.0001.

After 100 generations of directed Adaptive Laboratory Evolution (ALE) (32) of the mL50_1-249_ strain, mito-translation function was recovered through accumulation of mutations. Our ALE scheme selected for rapid growth of the mL50_1-249_ strain in non-permissive media over 100 generations (Figure 2c). After 100 generations, three colonies from three independent evolution cultures were isolated. Mito-translation was assayed in each in non-permissive media by fluorescence of mito-sfGFP. Mito-translation function in all three isolates appeared, by these measures, to be equivalent to MOY1355 (Figure 2c). An alternative evolution scheme of oscillating between permissive and non-permissive media was tested, but showed a less consistent increase in mito- ribosome function (Supplementary Figure 3).

#### Truncated Mito-rProtein uL23

uL23 is a universal rProtein located adjacent to the exit tunnel, with homologs across the tree of life. uL23 is essential for translation, and impacts nascent protein folding (33). As an architectural component of the diverted exit tunnel of the *S. cerevisiae* mito-ribosome (Figure 3a) (31), changes in mito-uL23 are expected to impact mito-translation.

**Figure 3.**
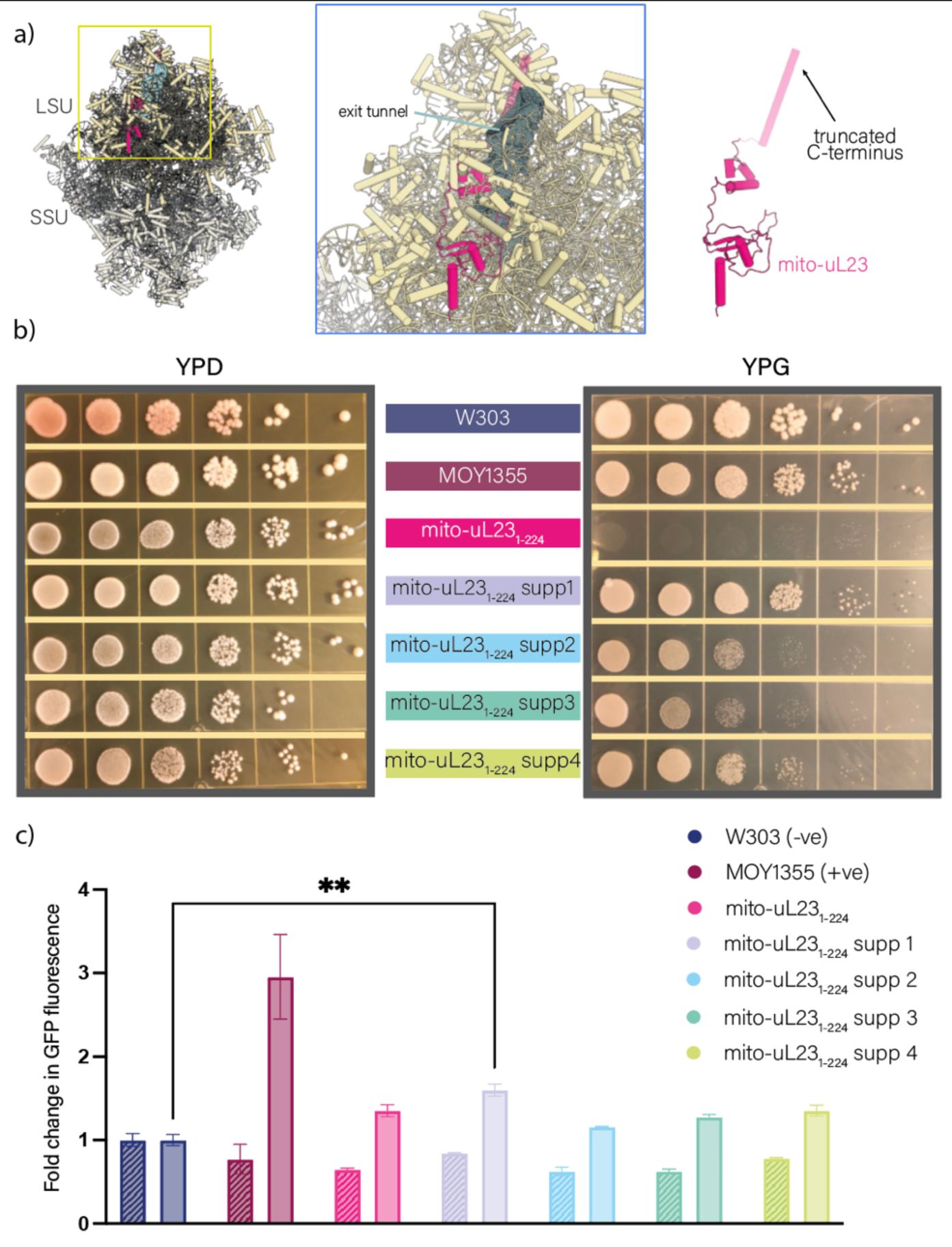
TS function can be recovered via ALE. a) mito-TS function was compromised by engineered truncation of mito-uL23 (to give mito-uL23_1-224_, red). The nascent peptide exit tunnel is indicated. b) Growth of wild-type, the mito-uL231-224 strain and four suppressors on both permissive (YPD) and non-permissive (YPG) media. c) Function of the mito-TS assayed in all strains via flow cytometry after growth in permissive (YPD) and non-permissive (YPG) media. Growth in permissive media is indicated with diagonal hashes. Growth in non-permissive media is indicated by solid fill. Mean fluorescence intensity values were divided over W303 for the same experiment and media conditions to determine fold change over background (N = 10,000). ns = p > 0.05, * = p ≤ 0.05, ** = p ≤ 0.01, *** = p ≤ 0.001, *** = p ≤ 0.0001.

We edited mito-uL23 by removing the C-terminal α-helix. Mito-uL23_1-261_ (1-261 indicates the length of the native mito-rProtein) was truncated in MOY1355 by the removal amino acids 225- 261 to create mito-uL23_1-224_. The results suggest that this edit shuts down mito-translation. No growth of the mito-uL23_1-224_ strain was observed on non-permissive media (Figure 3b). Fluorescence of mito-sfGFP in the mito-uL23_1-224_ strain measured via flow cytometry was equivalent to in the non-fluorescent control (Figure 3c).

We re-evolved mito-translation functionality in the mito-uL23_1-224_ strain. As with antibiotic resistance evolution, mutations were accumulated during growth in permissive media followed by selection in non-permissive media. Four suppressor strains of the mito-uL23_1-224_ phenotype, that grew on non-permissive media, were identified (Figure 3b). Sequencing revealed all four suppressors had a S39I change in protein Rrg9 and truncations in protein AI1 that were not present in mito-uL23_1-224_ pre-evolution strain (Figure 4a). While the function of Rrg9 is unknown, it localizes in the mitochondrion and interacts with components of the mito-translation system (Figure 4b) (34–37). Deletion of *RRG9* renders *S. cerevisiae* unable to respire (38). The mito-translation function of the four mito-uL23_1-224_ suppressors in non-permissive media remained low, with only Suppressor 1 having mito-translation function above W303 background fluorescence levels with a 1.6-fold increase. Recreation of the S39I SNP in Rrg9 yielded a petite positive phenotype and subsequent fluorescence levels similar to WT MOY1355. The addition of the mito-uL23_1-224_ truncation results in reduced mito-translation compared to Rrg9 S39I without the truncation, but respiration and mito-translation still occur indicating that the Rrg9 S39I SNP is solely responsible for recovery (Figure 5). AI1 is not known to interact with proteins involved in mito-translation (supplementary Figure 4).

**Figure 4.**
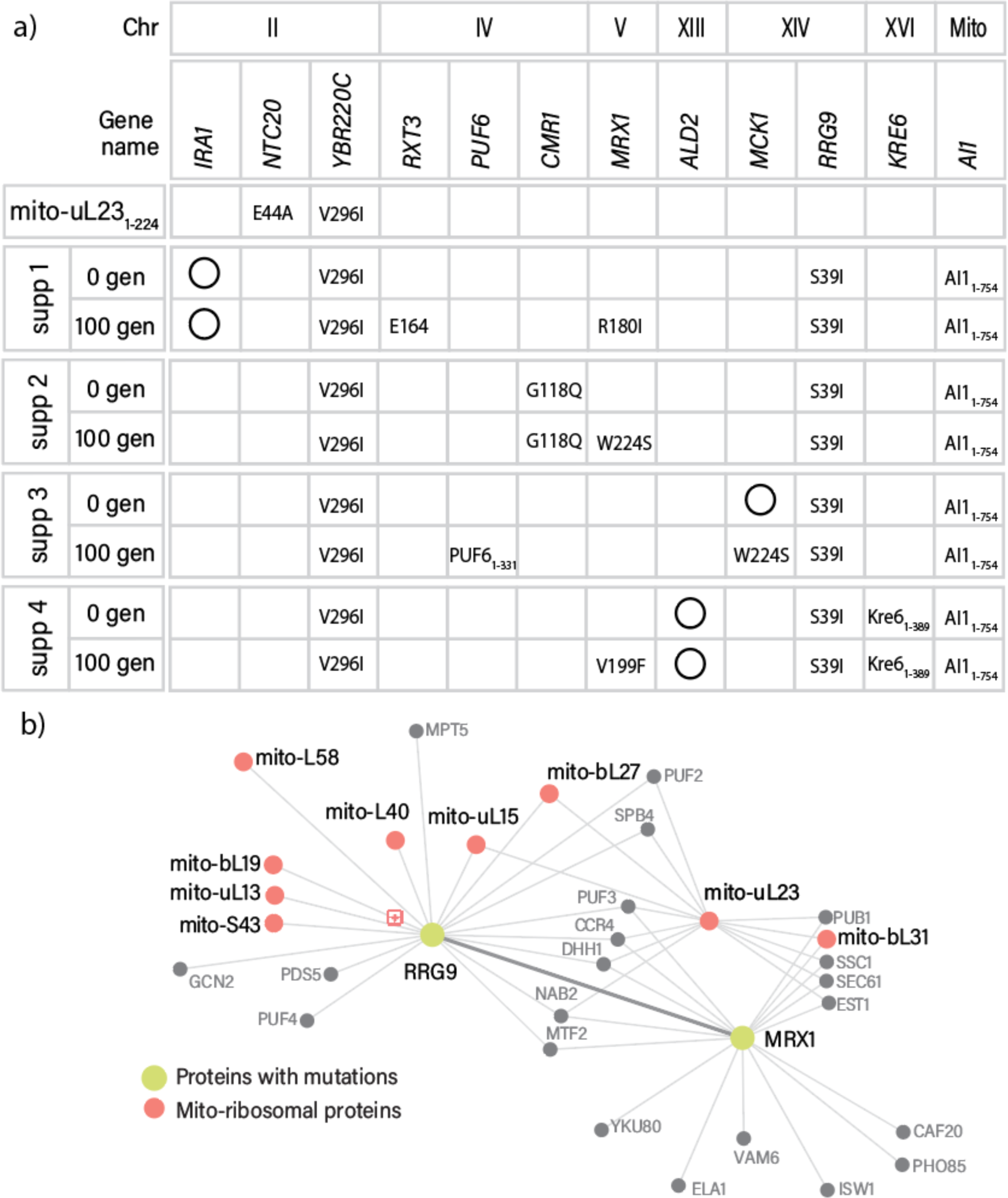
mito-uL23^1-224^ suppressors. a) Mutations found in protein-coding genes in mito-uL231-224 and four suppressors. 0 gen: initial suppressor. 100 gen: suppressor strain after 100 generations. Synonymous mutations (open circles), non-synonymous (amino acid changes), and truncations are indicated on the chart. b) Map of physical interactions of RRG9 and MRX1. Proteins are represented by nodes, and physical interactions curated by BioGRID (39) are represented by lines between nodes. The connection between RRG9 and MRX1 is highlighted by a bold line. Green nodes: mutated proteins in suppressor 4. Pink: ribosomal proteins and other proteins related to mitochondrial translation.

**Figure 5.**
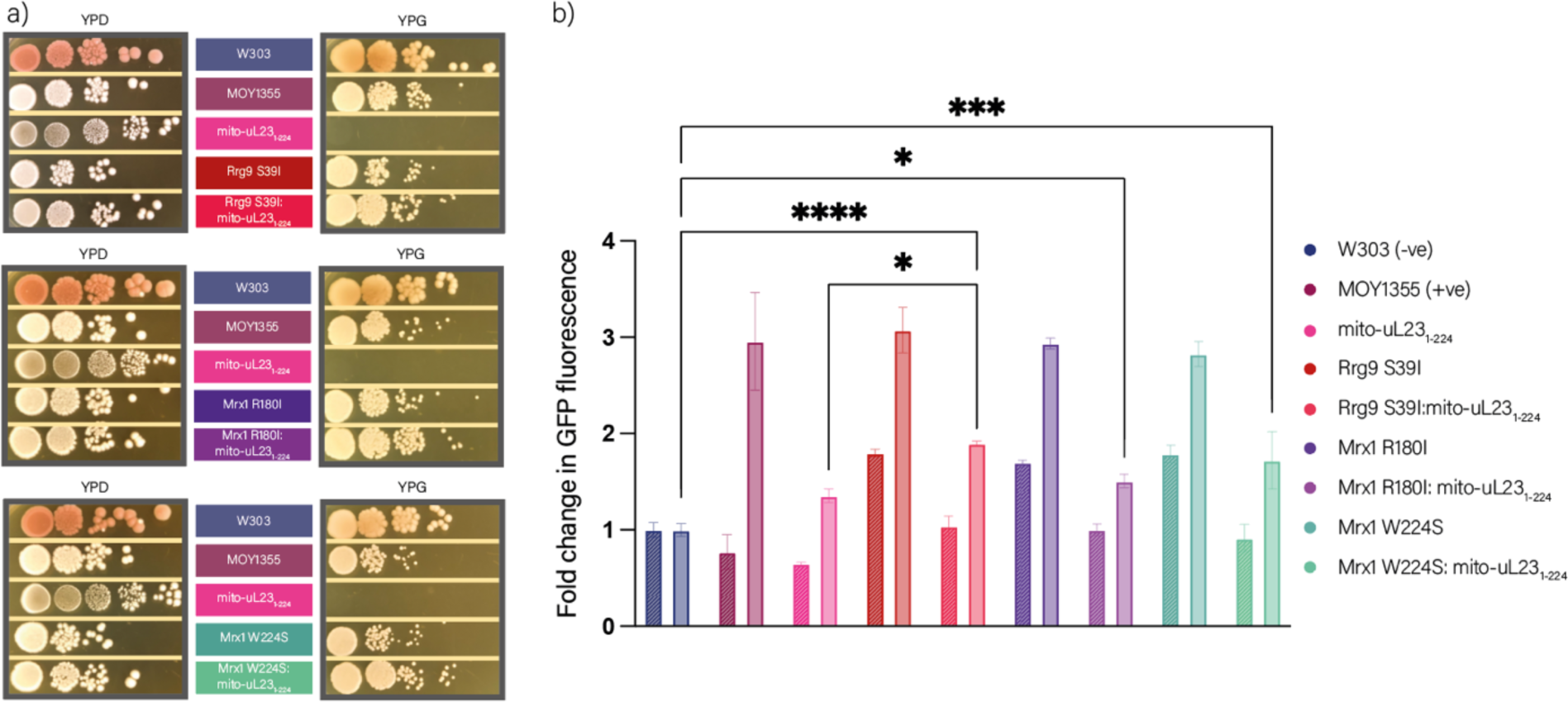
Engineered recreation of SNPs in mito-uL23_1-224_ suppressors and evolved strains. a) Growth of wild-type strains, mito- uL23_1-224_, and SNPs with and without mito-uL23 truncation on both permissive (YPD) and non-permissive (YPG) media. b) Function of the mito-translation system assayed in all strains in the flow cytometer after growth in permissive (YPD) and non-permissive (YPG) media. Permissive media is denoted with diagonal lines and non-permissive with solid fill. Mean fluorescence intensity values were divided over W303 for the same experiment and media conditions to determine fold change over background (N = 10,000). ns = p > 0.05, * = p ≤ 0.05, ** = p ≤ 0.01, *** = p ≤ 0.001, *** = p ≤ 0.0001.

The ALE method established with mL50_1-249_ was used for directed evolution of the mito-uL23_1-224_ strain. Over 100 generations, mito-translation function recovered through accumulation of mutations. Three colonies were isolated from each of two independently evolved cultures (per suppressor) and mito-translation was assayed in non-permissive media by growth rate on non- permissive media and by fluorescence of mito-sfGFP. Increased mito-translation function compared to the original suppressor was observed in most isolates (supplementary Figure 5). We qualitatively assayed mitochondrial TS using confocal microscopy to ensure increasing fluorescent values in evolved strains were due to an increase in mito-translation and not an increase in number of mitochondria. Images of MOY1355, mito-uL23_1-224_, and suppressors showed the number of mitochondria to be constant (supplementary Figure 6).

Three of the four suppressors evolved over 100 generations acquired mutations in protein Mrx1 (R180I, W224S, or V199F; Figure 4a). Mrx1 is known to associate with the mito-ribosome and other factors involved in mito-translation (Figure 4b). Deletions of *MRX1* are respiration deficient indicating it is essential for respiration. We specifically recreated the Mrx1 R180I and W224S SNPs and show they do not affect mito-translation in MOY1355 (Figure 5b). Both the Mrx1 R180I and W224S SNPs recover mito-translation above background levels in the mito-uL23_1-224_ strain (Figure 5b). Additional non-synonymous mutations were found in *MCK1*, *PUF6*, and *RTX3* (Figure 4a; Supplementary Figure 4).

### Replacement of mito-rProteins

The *S. cerevisiae* mito-translation system can be used as a platform for replacing ribosomal proteins with homologs from other species, and/or with ancestral reconstructions. We used rProtein uL2 to test the ability of yeast to survive and respond to global and local modifications to mito-uL2. rProtein uL2 is a universal ribosomal protein that is essential for peptidyl transfer (39). To measure the impact of a global but minimal change, we replaced one mito-uL2 gene (*RML2)* with another, while holding the amino acid sequence constant. We substituted the *S. cerevisiae* mito-uL2 sequence with a sequence optimized for cytosolic expression in *S. cerevisiae.* The resulting altered *RML2* gene encoded the native mitochondrial amino acid sequence with 78% nucleotide identity to the WT gene (supplementary Table 2). The resulting strain (mito- uL2_codon-optimized_) grew more slowly than wild-type in non-permissive media (Figure 6c) and had slightly reduced mito-ribosomal function compared to MOY1355 when assayed by mito-sfGFP, from 3.0-fold over background to 2.5-fold over background (Figure 6d).

**Figure 6.**
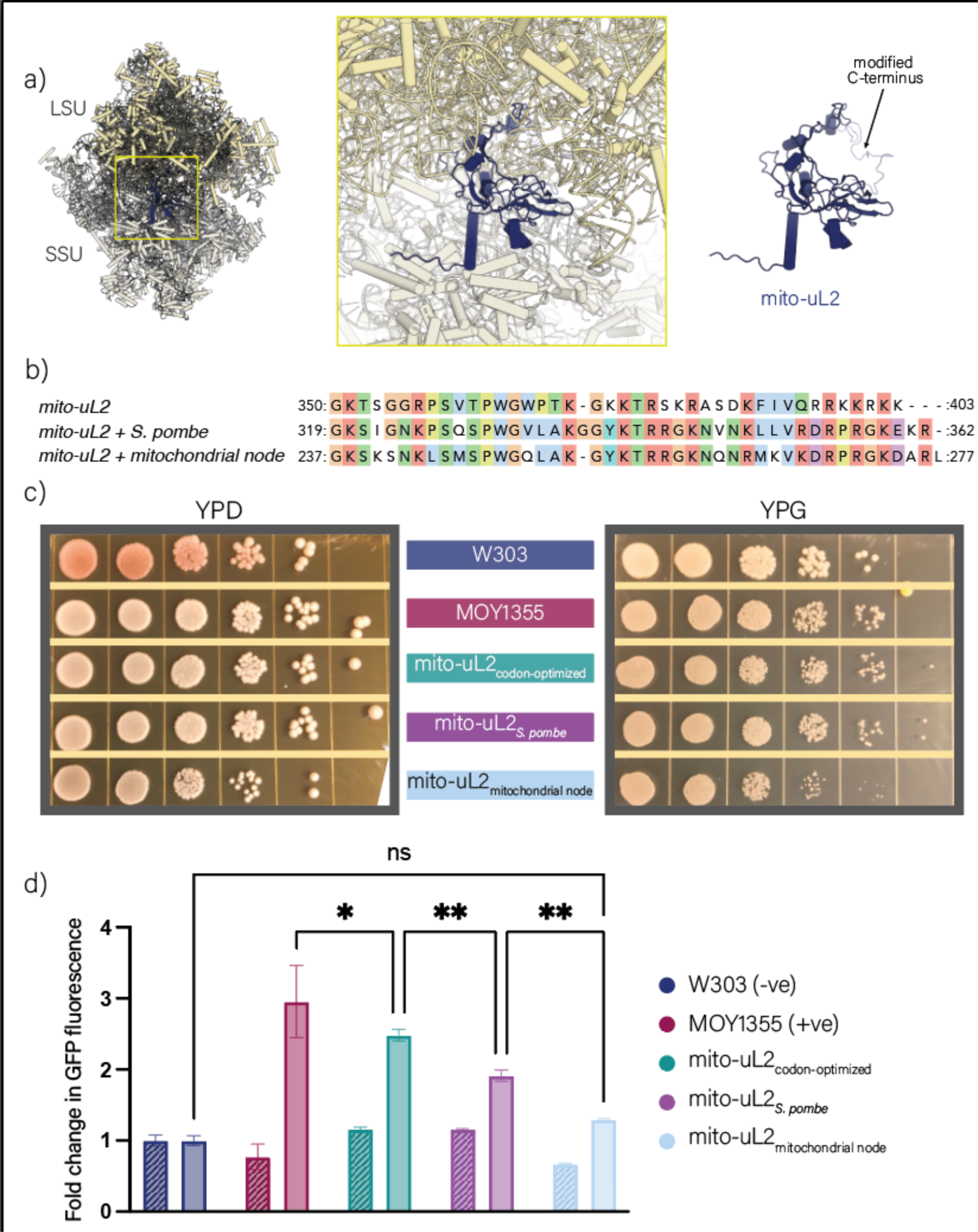
Modifications to mito-uL2 decrease mito-translation. Location of the mito-uL2 truncation in the mito-ribosome (PDB code 5MRC). a) Location of the mito-uL2 truncation in the mito-ribosome (PDB code 5MRC). mito-uL2 is shown in blue. b) Multiple sequence alignment of the 44 C-terminal amino acids of mito-uL2 in *S. cerevisiae* (RefSeq NP_010864.3)*, Schizosaccharomyces pombe*(RefSeq NP_587925.2) and the inferred ancestral reconstruction of the mitochondrial common ancestor. Residues are colored using the Clustal X color scheme. c) Growth of wild-type strains, mito-uL2_codon-optimized_, mito-uL2*_S. pombe_*, and mito- uL2_mitochondrial node_ in both permissive (YPD) and non-permissive (YPG) media. d) Function of the mito-translation system assayed in all strains in the flow cytometer after growth in permissive (YPD) and non-permissive (YPG) media. Permissive media is denoted with diagonal lines and non-permissive with solid fill. Mean fluorescence intensity values were divided over W303 for the same experiment and media conditions to determine fold change over background (N = 10,000). ns = p > 0.05, * = p ≤ 0.05, ** = p ≤ 0.01, *** = p ≤ 0.001, *** = p ≤ 0.0001.

Comparisons of experimental structures of mito-ribosomes reveal that the C-terminus of mito- uL2 is structurally variable over phylogeny, making this region of mito-uL2 an interesting target for perturbation. We replaced 44 C-terminal amino acids of *S. cerevisiae* mito-uL2 with 44 C- terminal amino acids of the *Schizosaccharomyces pombe* homolog of mito-uL2. This substitution changed 29 of 44 amino acids of the C-terminal tail (Figure 6b). The resulting growth was comparable in non-permissive media to the codon-optimized mito-uL2, although mito-translation function decreased from 2.5-fold to 1.9-fold over background.

Lastly, we inferred the amino acid sequence of an ancestral mito-uL2 from a multiple-sequence alignment of uL2 and mito-uL2 homologs across the tree of life. We replaced the 44 C-terminal amino acids of the *S. cerevisiae* mito-uL2 with the C-terminus of the inferred ancestral reconstruction of the mitochondrial common ancestor with a total of 27 amino acids changed and 3 deleted (mito-uL2_mitochondrial node_; Figure 6b). In non-permissive conditions, this modification reduced growth (Figure 6c). Mito-uL2_mitochondrial node_ could grow on non-permissive media, although mito-translation is reduced (Figure 6d). These results are consistent with the divergence in amino acid properties in the replaced segment of mito-uL2 (Figure 6b).

## Discussion

Nature offers us a manipulable, evolvable, fully orthogonal TS in the mitochondrion of *S. cerevisiae*. The primary TS is constrained to the cytosol and is descended from the TS of an archaeon (40). The orthogonal TS is in a mitochondrial box, and is descended from the TS of an ⍺-proteobacterium (41,42). Genes encoding mito-translation factors, mito-aminoacyl-tRNA synthetases - (aaRS’s) and most mito-rProteins are located in the nuclear genome. These proteins are directed to mitochondria by targeting sequences (mitochondrial localization signals) (43). Genes encoding mito-rRNAs, mito-tRNAs and one mito-rProtein (mito-uS3) are located in the mitochondrial genome, which also encodes seven integral membrane proteins required for respiration (43). The primary *S. cerevisiae* TS is necessary for survival while the orthogonal TS is conditionally necessary for survival.

The complete orthogonality of the mito-TS of *S. cerevisiae*, and the conditional dependence for survival on its function, offers the possibility of continuous, simultaneous directed evolution of all TS components. The combined results here indicate that perturbations of the mito-TS can cause repeatable and generally interpretable compensatory changes to mito-TS components over evolution.

The functionality of the mito-TS was quantitated here via flow cytometry, growth on non-permissive media and sequencing. The fluorescence intensity of mito-sfGFP was used as a proxy for functionality of the mito-TS. Microscopy results here are consistent with a previous demonstration of containment of mito-sfGFP within mitochondria and the lack of change in the number of mitochondria over the course of our experiments (Supplementary Figure 5). Fluorescence of mito-sfGFP indicates that the unusual path of the *S. cerevisiae* mito-TS tunnel (29) does not prevent folding of soluble proteins.

We validated the general utility of the *S. cerevisiae* mito-TS in directed evolution using well-characterized mito-ribosome-targeting inhibitors, chloramphenicol and erythromycin. The SNPs that conferred resistance to these antibiotics had been previously identified (27,28) but resulting mito-translation had not been quantified. Sequencing, flow cytometry and growth on non-permissive media indicate full recovery of TS function is achieved by mutations of the LSU mito- rRNA.

We investigated the evolutionary response of the mito-TS to changes in mito-rProteins. We re-evolved full mito-TS functionality after truncations of mL50 or mito-uL23. Suppressors of mito- uL23 truncation phenotype, with recovered function, consistently show mutations in mito- proteins Rrg9 and Mrx1. These proteins are known to interact with the mito-ribosome (37) and cause respiratory deficiency when absent (38,44,45) but have not previously been ascribed function. The results here do not define the function of Rrg9 or Mrx1 but allow to propose that Rrg9 and Mrx1 interact with the mito-TS and have critical role in its function. The evolutionary method developed here appears to be a general approach for discovering functional dependencies. Recreations of the SNPs identified in the WGS revealed that they do recover some level of mito-translation when combined with the mito-uL23 truncation.

We probed the effect of replacement of universal rProtein mito-uL2 on the mito-TS. We substituted the native *RML2* gene with a codon-optimized sequence and then further replaced the C-terminal tail with sequences from *S. pombe* and the hypothesized sequence from the mitochondrion in the last universal eukaryotic ancestor. To our knowledge *in vivo* ancestral reconstructions of parts of the ribosome have not been attempted previously. These modifications showed reduced function and could be further evolved as with mito-uL23 to start to push the ribosome back in time.

There are certain disadvantages to directed evolution of the mito-TS. mito-rRNAs and mito-tRNAs are encoded on the mito-genome, and are difficult to manipulate by conventional techniques of molecular biology. The mitochondrial genome can be manipulated using biolistic bombardment and homologous recombination (46). Less precise approaches to mito-genome engineering use base editors, like deaminases, fused to mitochondrial targeting signals (47). A second disadvantage to the mito-TS is the propensity for *S. cerevisiae* become petite, i.e., to lose mito- genomes when cells lack specific proteins associated with mitochondrial transcription and translation (e.g. mL54, mS35, Mtf1) (38,48). We showed the upregulation of *RNR1* decreased the petite formation rate, even with a strain with lower mito-translation functionality (Supplementary Figure 7).

The mito-TS can be used for incorporation of non-canonical amino acids. The *E. coli* tyrosine aaRS, when fused to a targeting tag, has been shown to specifically charge the *S. cerevisiae* mitochondrial tRNA^Tyr^. Alterations to the *E. coli* aaRS^Tyr^ have been shown to incorporate photoaffinity probes, bioconjugation handles, and a nonhydrolyzable mimic of phosphotyrosine into proteins (49). Furthermore, there are unused codons in the mito-genome, including the CGN family of arginine codons (50), offering an opportunity for genetic code expansion that will not affect native mito-translation.

Full control of the TS will allow expansion of the sidechain inventory of biopolymers and transformation of the backbone into new chemical spaces. These advances will require that we overcome evolutionary forces that have held the TS essentially fixed for billions of years. Advances require transformation of catalytic competencies of the PTC and aaRSs and of delivery/transport functionality of the exit tunnel (33,51–53). A host of factors including rRNAs, tRNAs, translation factors, aaRS’s, and rProteins are directly involved in these processes and many other factors are indirectly involved. Continuous and comprehensive directed-evolution of a fully orthogonal TS *in vivo* appears to be the most direct for technical control of the TS. The work here indicates the power of system-wide evolutionary approaches for translation system engineering.

## Materials and Methods

### Strains, plasmids and oligonucleotides

The W303 strain was a gift from Dr. Thomas Petes (Duke University). The MOY1355 strain (Mat α ade2-1 his3-11,15 trp1-1 leu2-3,112 ura3-1, sfGFPm:cox2, COX2) (23) was a gift from Dr. Martin Ott (Stockholm University). Other strains were derivatives of MOY1355. Freshly thawed yeast strains (if capable of respiration) were streaked onto YPG or (if incapable of respiration) YPD and grown at 30 °C in the presence of atmospheric O_2_. Permissive media (YPD) is 1% yeast extract, 2% peptone, 2% glucose and non-permissive (YPG) is 1% yeast extract, 2% peptone, 3% [v/v] glycerol). Uracil dropout, YPD + G418 (0.3 mg/mL) and 5-fluoroorotic acid were used for the development of strains. Strains were constructed either with the *kanMX* marker (mL50 and uL23 perturbations) or CRISPR (uL2 replacements). *kanMX* and *URA3* marker truncations were made by amplifying the *kanMX* region of pKL55 or *URA3* of pFL34 using primers with 40 bp homology to the gene sequence for *mrp20* and *mrpl13*. The uL2 strains and SNP strains were created using CRISPR/Cas9 to replace the existing *RML2* gene with a synthesized gene.

### CRISPR

gRNAs were designed using the CRISPR Design Tool in Benchling and assembled into the CRISPR backbone (Addgene plasmid #198866) using Golden Gate assembly. The assembled plasmids were transformed into *E. coli* dh5⍺ and grown overnight on LB + 50 µg/mL spectinomycin at 37 °C. Non-fluorescent colonies were picked, cultured overnight in liquid LB + 50 µg/mL spectinomycin and miniprepped using the Qiagen miniprep kit (Cat. No. / ID: 27104). Plasmid assembly was verified by sequencing (Eurofins Genomics).

### Yeast Transformation

To select for cells capable of respiration, cells were inoculated in 5 mL liquid YPG for the initial growth from colonies for 2 days, shaking at 200 RPM. The starter culture was back-diluted in 50 mL YPD to a final OD of ∼0.15 and returned to the incubator. When the culture reached an OD of 0.6-0.8 the cells were pelleted by centrifugation at 1363xg for 5 min (Sorvall Legend XTR centrifuge with a TX-750 Swinging Bucket Rotor, ThermoFisher). The supernatant was decanted and the pellet resuspended in 20 mL autoclaved H_2_O and pelleted again. The supernatant was discarded and the pellet resuspended in 400 µL H_2_O. 150 µL of cell suspension was transferred to a 1.5 mL Eppendorf tube and spun for 30 seconds at 13,000 rpm in a tabletop centrifuge. The supernatant was discarded and the pellet was resuspended in 148 µL DNA mixture (for CRISPR: 200 ng CRISPR plasmid, 1000 ng donor DNA (NotI digest plasmid); for *kanMX*: 40µL gel-purified DNA, 108 µL H_2_O) and 20 µL salmon sperm DNA (boiled for 5 min, chilled on ice for 3 min). PEG (50%, MW 3350; 480 µL) then 72 µL 1M LiAC were added. The mixture was vortexed and incubated at 30 °C for 30 mins, incubated at 42 °C for 15 mins then plated on uracil dropout (6.8 g/L Yeast Nitrogen Base Without Amino acids (Sigma, Catalog number: Y0626), 2% glucose, 1.92 g/L amino acid mix without uracil (Sigma, Catalog number: Y1501) or on YPD. Ura+ transformants were grown for three days at 30 °C. To select for drug resistance, cells were grown on YPD for one day and replica-plated onto YPD + G418 (0.3 mg/mL) plates. Colonies were picked and insertions were verified using colony PCR with Phire Plant Direct PCR Master Mix (ThermoScientific, Catalog number: F160L) followed by sequence verification (Eurofins Genomics).

### Evolution

Yeast strains were evolved via spontaneous mutation accumulation for both antibiotic resistance and suppression of the YPG- phenotype of the uL23 truncation. Mutations were accumulated inoculating colonies in 5 mL YPD and incubating overnight with shaking at 200 RPM. The saturated culture was plated on 20 plates (200 µL per plate) of either YPG, YPG + erythromycin (2 mg/mL) or YPG + chloramphenicol (2.5 mg/mL). The plates were monitored for 2 weeks of incubation and antibiotic resistant strains and suppressors were picked. For strains exhibiting antibiotic resistance, rRNA was sequenced first (Eurofins Genomics) followed by whole genome sequencing (Psomagen).

The Adaptive Laboratory Evolution protocol was adapted from Espinosa, *et al.* (54). Colonies were picked from YPG plates and grown overnight in liquid YPG, shaking at 200 RPM. After recording the OD, the culture was back-diluted to OD 0.1 in YPG or YPD, depending on the evolution scheme. The OD measurement and back-dilution process was repeated daily in either YPG only or oscillating between YPD and YPG until about 100 generations. The final culture was diluted in autoclaved H_2_O and plated on YPG. Three colonies were picked from each evolution culture for further analysis.

### Assay of Petite Formation

Three colonies per strain were picked from YPG plates and each was used to inoculate 5 mL liquid YPG. The cultures were grown at 30 °C and shaking at 200 RPM until saturated (2 days). Cultures were back-diluted in 5 mL YPD to an OD of ∼0.003. The diluted cultures were once again grown to full density over 24 hours at 30 °C. Final OD measurements revealed ∼12 generations of growth in YPD (within the recommended 10-15 generation range) (55). The OD was recorded and the culture was diluted in autoclaved H_2_O and plated on YPD (∼100 colonies per plate, 3 plates per culture). The plates were transferred to a 30 °C incubator for 2 days. Colonies were counted and replica-plated onto YPG. The YPG plates were incubated a 30 °C for 1 day. Colonies that grew on YPG maintained the ability to respire and counted as non-petite.

### In silico

#### Homology searching

The multiple-sequence alignment (MSA) of uL2 sequences from 66 bacteria and 52 archaea representing a Sparse and Efficient Representation of Extant Biology (8) was retrieved from ProteoVision (56). Mitochondrial sequences of uL2m from five Jakobid species (57) (*Jakoba libera*, *Andalucia godoyi*, *Seculamonas ecuadoriensis*, *Jakoba bahamiensis*, and *Reclinomonas americana*) were added to the MSA of uL2 with MAFFT v7.490 (58). The final MSA containing archaeal, bacterial, and mitochondrial sequences that displayed a block structure with universally conserved, archaea-specific, and bacteria/mitochondrial-specific blocks. The MSA was trimmed to the universal blocks with BMGE v1.12 (59) with default parameters.

### Ancestral sequence reconstruction

A maximum likelihood tree of the trimmed MSA was calculated with PhyML 3.0 (60) on the Montpelier bioinformatics platform. The substitution model (LG + G + I) selection was made with SMS (61) using a Bayesian information criterion. The archaeal branch was used as an outgroup to root the ML tree. The joint reconstructions of all ancestral nodes were calculated with the GRASP (Graphical representation of ancestral sequence predictions) web server (62) using the Jones-Taylor-Thornton (JTT) evolutionary model.

The ThermoFisher GeneOptimizer tool was used to codon-optimize the amino acid sequences for cytosolic expression in *S. cerevisiae.* The resulting DNA sequences were synthesized by GeneWiz in the pUC-GW-Kan vector and then assembled into the *RML2* ORF CRISPR backbone with 510 bp homology arms using Golden Gate assembly designed in Benchling. The assembled plasmids were transformed into *E. coli* dh5⍺ and grown overnight on LB + 25 µg/mL chloramphenicol at 37 °C. Non-fluorescent colonies were picked and grown overnight in liquid LB + 25 µg/mL chloramphenicol and miniprepped using the Qiagen miniprep kit (Cat. No. / ID: 27104). Plasmid assembly was verified by sequencing (eurofins Genomics).

### Growth Assays

To monitor the growth rate a colony was diluted in 200 µL H_2_O. This mixture then went through a serial dilution of 30 µL in 170 µL of H_2_O five times. 10 µL of each dilution was spotted on YPD and YPG (with the addition of antibiotics for the Cam^R^ and Ery^R^ experiments). Growth at 30 °C was monitored over four days. Photographs were taken to observe and record colony size.

### Flow cytometry

Three colonies per strain were picked and inoculated in filtered YPD + 40 mg/L adenine hemisulfate and grown overnight. The cultures were back-diluted 1:100 in 5 mL fresh YPD and harvested after 24 hours growth at 200 rpm for flow cytometry. The strains were further diluted 1:50 in 5mL fresh YPG and grown for a further 24 hours before harvesting for flow cytometry.

Samples were diluted 1:100 in 200 µL H_2_O in a 96 well plate for flow cytometry. The samples analyzed with a a Cytoflex S flow cytometer with 10,000 samples recorded per well (FSC: 86, SSC: 119, FITC: 149). The collected FITC, SSC and FSC data were analyzed using FlowJo, with doublets excluded via gating FSC-H plotted against FSC-A. The FITC values of the subsequent singlet populations were averaged per well to obtain fluorescence per cell. Analysis was conducted in Prism9 using one-way or two-way ANOVAs with multiple comparisons, depending on the data. P-values are denoted with ns = p >0.05, * = p ≤0.05, ** = p ≤0.01, *** = p ≤0.001, *** = p ≤0.0001.

### Confocal Microscopy

Colonies were picked and grown overnight in 5 mL YPG. One mL of culture was harvested by spinning at 13,000 rpm for 30 seconds. The supernatant was discarded and the pellet resuspended in 10-1000 µL H_2_O (depending on culture density). Two µL of each sample was pipetted onto a microscope slide (Electron Microscopy Sciences 63422-06) that had been prepared with 1.4% agarose.

Images were acquired by Zeiss LSM 700 laser scanning confocal microscope using Plan-Apochromat 63x/1.4 Oil lens with pinhole size set to 1AU. Line average of 4 was applied. sfGFP image was captured in EGFP setting with 488nm lasers set to 50% intensity, 500 gain, and 1.58µs pixel dwell across the different samples. Post imaging adjustment was performed in Zen 2012 software (Zeiss).

### Whole Genome Sequencing and Physical Interactions

Whole- genome sequencing was performed by Psomagen (Rockville, MD) using Illumina TruSeq DNA Nano 350 bp library prep and an Illumina NovaSeq 6000. Reads were aligned to the *S*. *cerevisiae* reference genome (yeastgenome.org) using the Burrows-Wheeler Aligner (63), Picard (http://broadinstitute.github.io/picard), SAMtools (64), and Genome Analysis Toolkit v3.8 (65). Variants were called against the respective reference genomes (MOY1355 for initial suppressors of antibiotic resistance and mito-uL23_1-224_ suppressors 1-4, Gen 0 for post-100 generation ALE) using Mutect2 (65). Physical interaction data was retrieved from BioGRID (66) for the proteins presenting mutations in mito-uL23_1-224_ and suppressors 1 to 4. The physical interactions between mutated proteins and other proteins were visualized with CLANS (67).

---

**Supplementary Figure 1.**
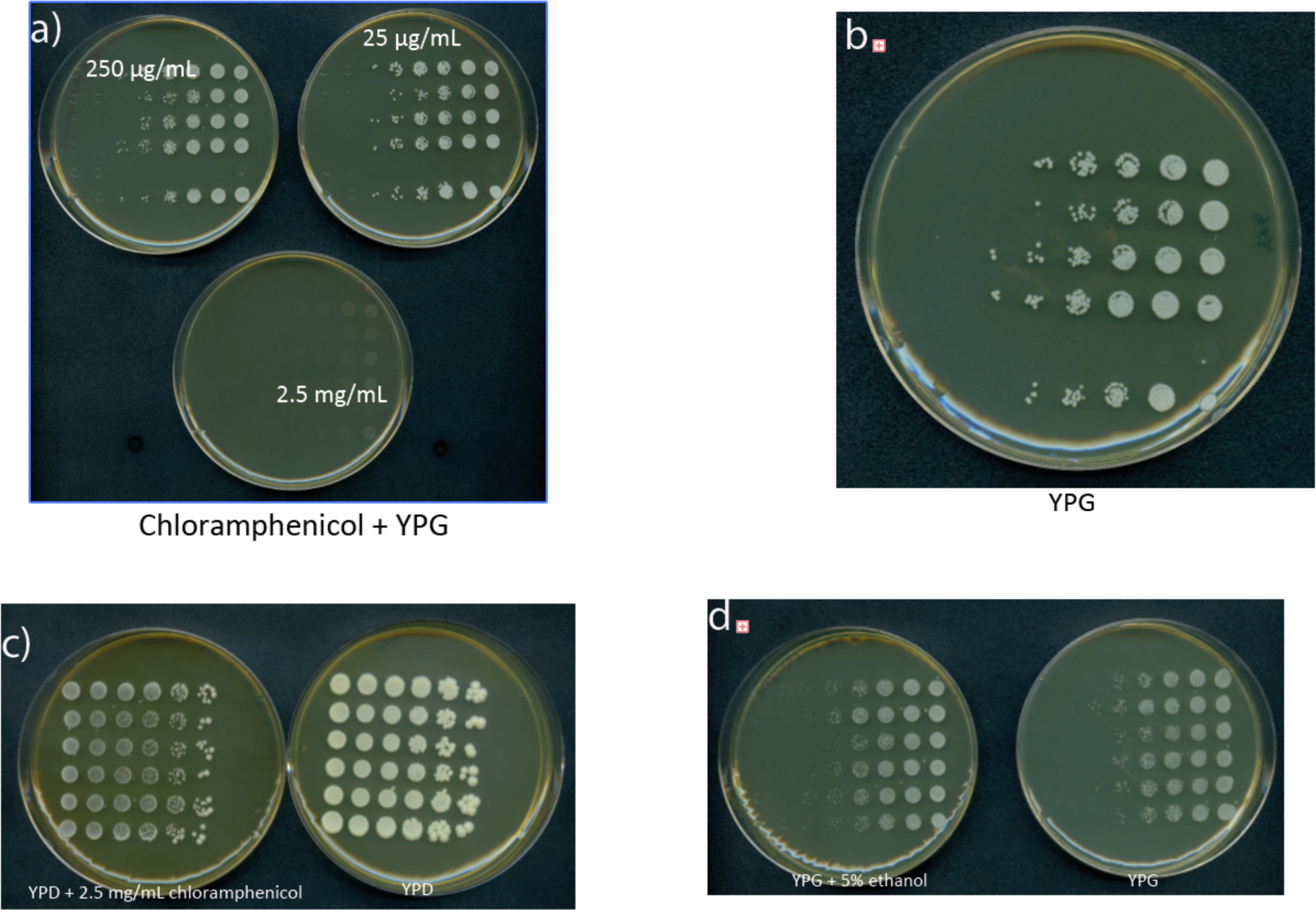
Growth of WT (MOY1355) on different carbon sources. a) The same strains on non-permissive media (YPG) with three concentrations of chloramphenicol. b) From the same experiment as a) but on non-permissive media (YPG) without antibiotics to indicate functionality of the mito-translation system. c) Growth on permissive media (YPD) with and without the effective dose of 2.5 mg/mL chloramphenicol. d) Growth on non-permissive media (YPG) with and without the addition of 5% EtOH, the ethanol concentration used with the chloramphenicol to indicate if that affects growth.

