## Supplementary Figures for "Translation in a Box: Orthogonal Evolution in the *Saccharomyces cerevisiae* Mitochondrion"

**Preprint Servers:** This manuscript was deposited as a preprint to bioRxiv under a CC-BY-NC-ND 4.0 International license.

**Classification:** Biological Sciences. Evolution.

**Keywords:** Orthogonal ribosome, translation

Brooke Rothschild-Mancinelli: orcid.org/0000-0003-0500-5903
Claudia Alvarez-Carreño: orcid.org/0000-0002-1827-8946
Loren Dean Williams: orcid.org/0000-0002-7215-4194

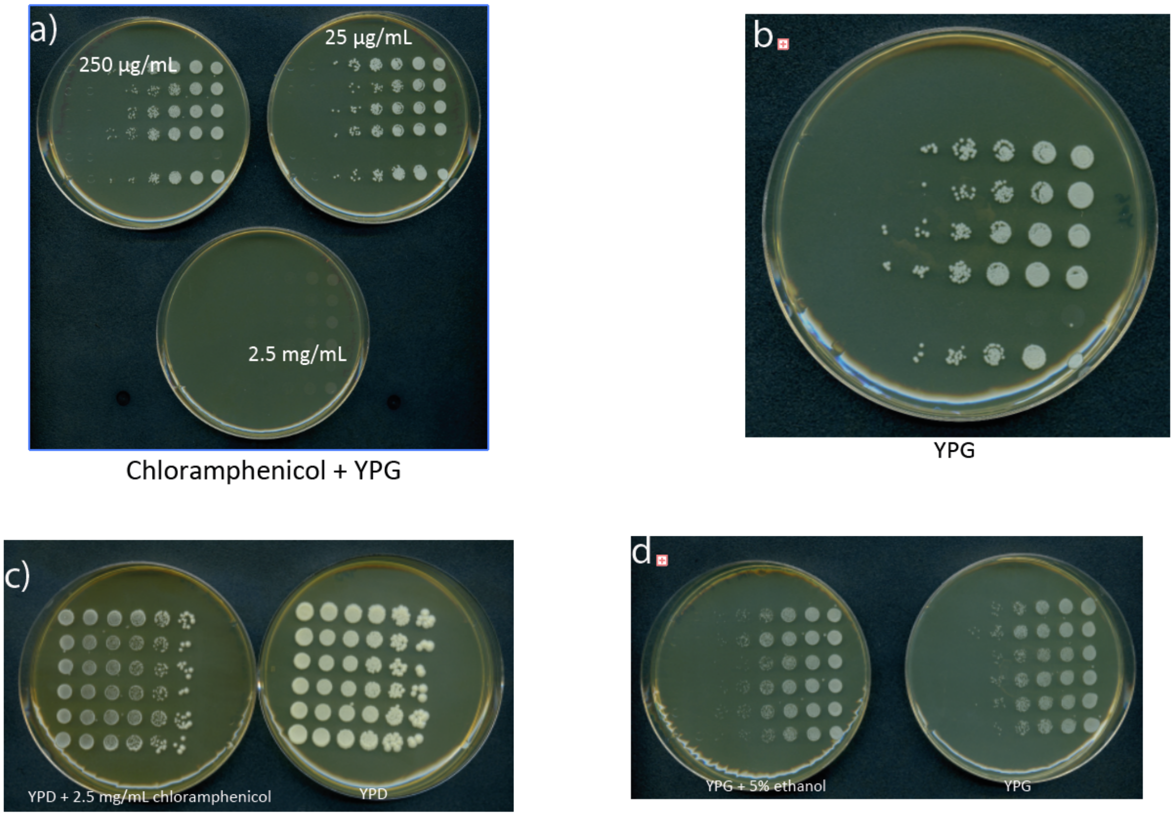


Supplementary Figure 1. Growth of WT (MOY1355) on different carbon sources. a) The same strains on non-permissive media (YPG) with three concentrations of chloramphenicol. b) From the same experiment as a) but on non-permissive media (YPG) without antibiotics to indicate functionality of the mito-translation system. c) Growth on permissive media (YPD) with and without the effective dose of 2.5 mg/mL chloramphenicol. d) Growth on non-permissive media (YPG) with and without the addition of 5% EtOH, the ethanol concentration used with the chloramphenicol to indicate if that affects growth.

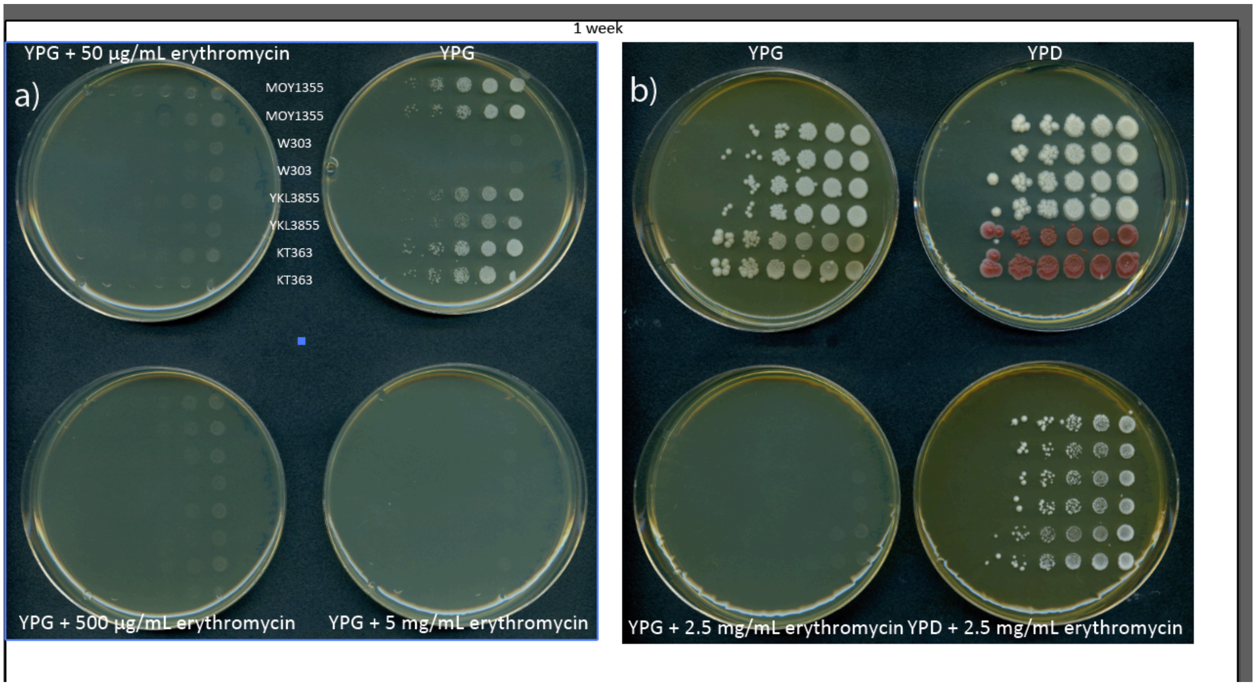


Supplementary Figure 2. Growth of WT (MOY1355) on different carbon sources. a) The same strains on non-permissive media (YPG) with three concentrations of erythromycin. b) From the same experiment as a) but on permissive media (YPD) with and without antibiotics to indicate functionality of the cytosolic translation system with erythromycin.

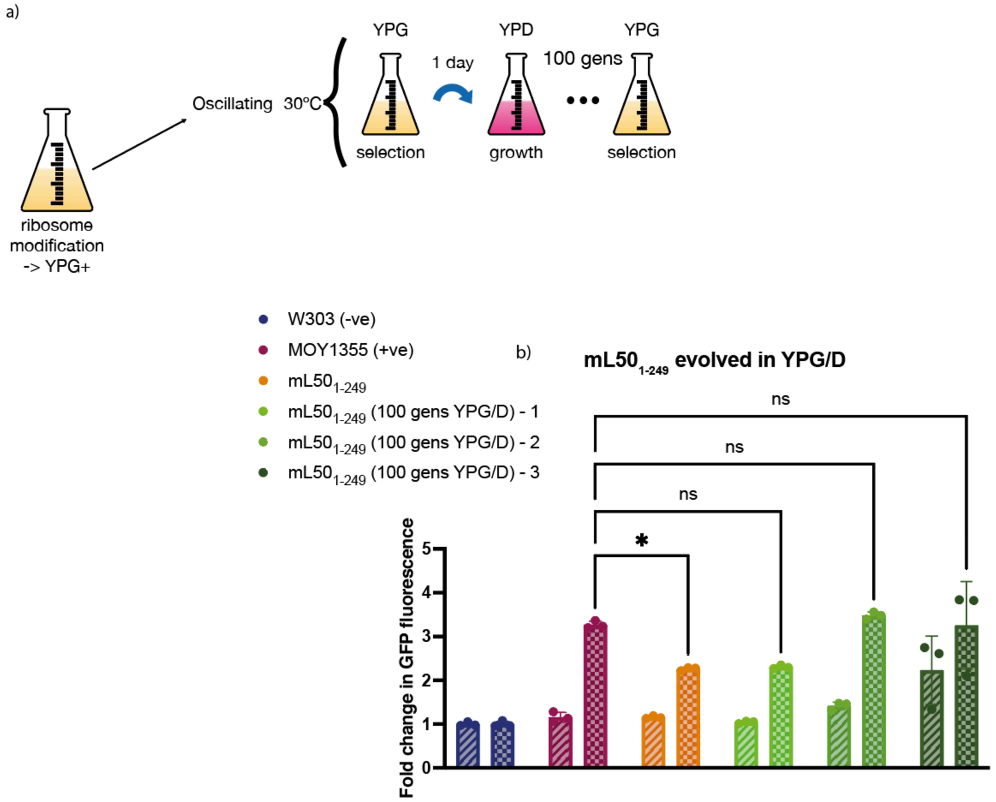


Supplementary Figure 3. Decreased function of mL50$_{1\text{-}249}$ can be recovered via ALE. a) the ALE evolution scheme used to increase mito-translation function oscillating between permissive (YPD) and non-permissive media (YPG). b) Function of the mito-translation system assayed on the flow cytometer after growth in permissive (YPD; diagonal stripes) and non-permissive (YPG; checked pattern) media. W303 (blue) is the non-fluorescent negative control, MOY1355 (maroon) is the WT fluorescent positive control derived from W303 and mL50$_{1\text{-}249}$ (orange) is the initial truncated strain with its evolved counterparts in shades of green. Flow cytometer values were divided over W303 for the same experiment and media conditions to determine fold change over background (N = 10,000).

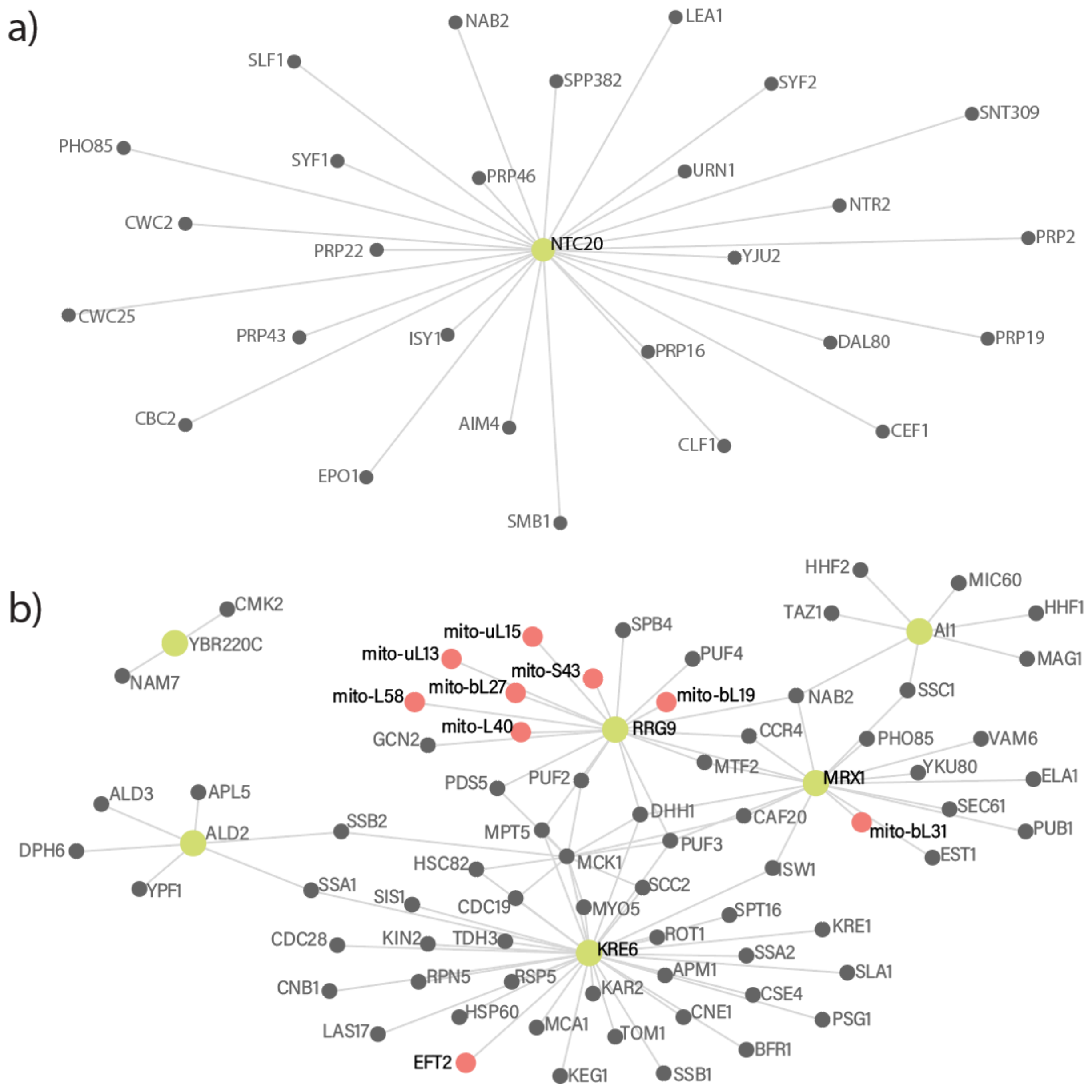


Supplementary Figure 4. a) Map of protein interactions of NTC20. b) Map of protein interactions of the proteins presenting mutations in suppressor 4: YBR220C, ALD2, RRG9, Kre6, and AI1.

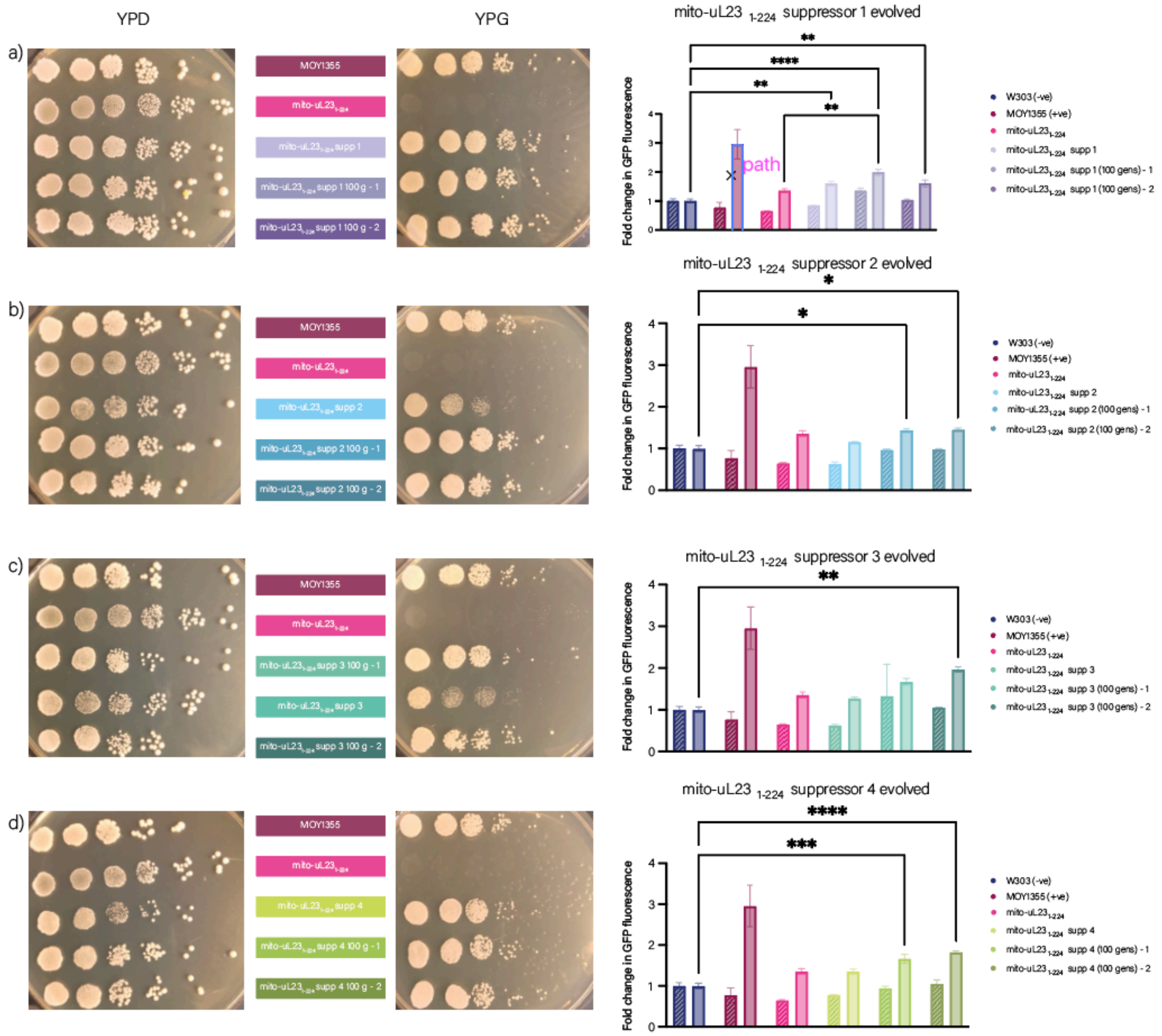


Supplementary Figure 5. Evolved Suppressors of mito-uL23$_{1\text{-}224}$ increase mito-translation function. a-c) Show the growth of the suppressors and their evolved strains on permissive (YPD) and non-permissive (YPG) media. The graphs show function of the mito-translation system assayed on the flow cytometer after growth in permissive (YPD; diagonal stripes) and non-permissive (YPG; checked pattern) media. W303 (blue) is the non-fluorescent negative control, MOY1355 (maroon) is the WT fluorescent positive control derived from W303. Mean fluorescence intensity values were divided over W303 for the same experiment and media conditions to determine fold change over background (N = 10,000).

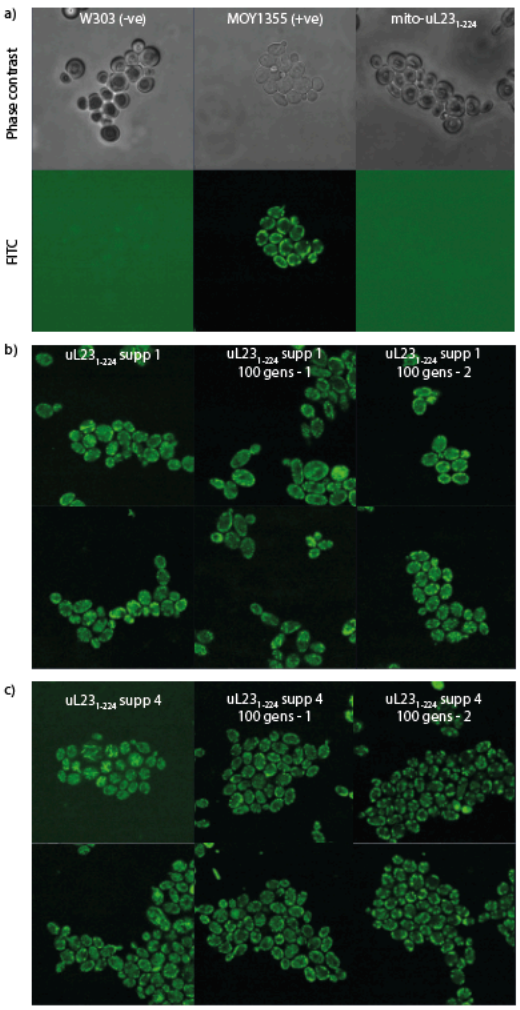


Supplementary Figure 6. Confocal microscopy of strains grown in non-permissive media. a) The control strains W303 (non-fluorescent), MOY1355 (fluorescent), and mito-uL23$_{1\text{-}224}$ (lacking mito-translation function) shown under phase contrast and the FITC channel. b) Suppressor 1 of uL23$_{1\text{-}224}$ with recovery of fluorescence and the two evolved strains from it. c) Suppressor 4 of uL23$_{1\text{-}224}$ with recovery of fluorescence and the two strains evolved from it.

The truncation of mL50 increased the petite formation rate of $mL50_{1-249}$ when compared to WT (MOY1355). The upregulation of *RNR1* has been shown to decrease petite formation frequency in wild type *S. cerevisiae* (1). The insertion of the strong promoter for *TDH3* upstream of *RNR1* rescues the petite formation rate of $mL50_{1-249}$ back to that of MOY1355 (supplementary Figure 4).

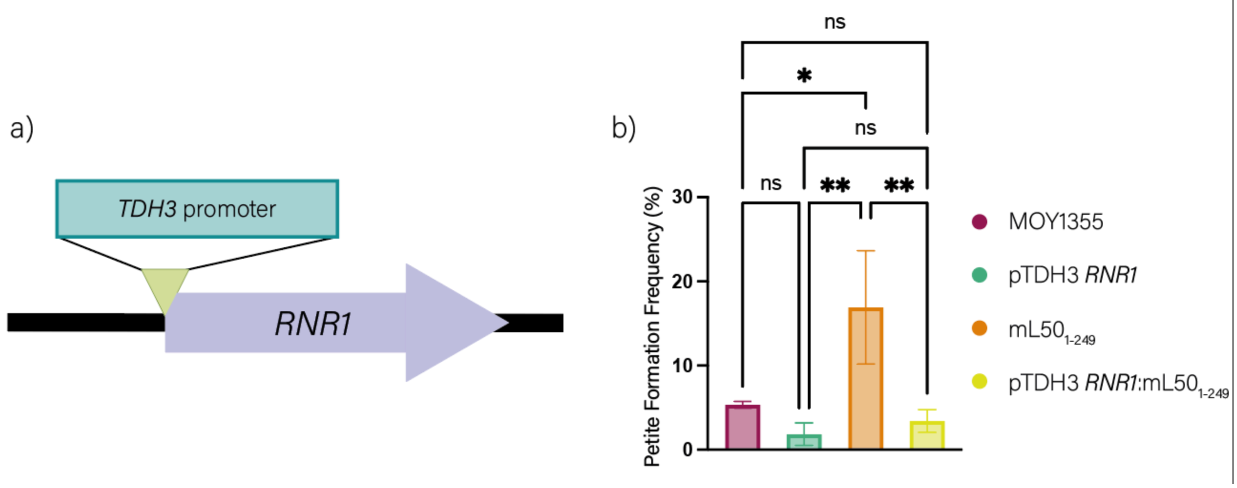


Supplementary Figure 7. Overexpression of *RNR1* decreases petite formation frequency. a) The precise insertion of the TDH3 promoter at the -1 position of the *RNR1* gene. b) MOY1355 (maroon), MOY1355 with the overexpression of *RNR1* (green), $mL50_{1-249}$ (orange), and $mL50_{1-249}$ with the overexpression of RNR1 (yellow). Analysis was done using an ordinary one-way ANOVA. ns = p >0.05, * = p ≤0.05, ** = p ≤0.01, *** = p ≤0.001, *** = p ≤0.0001.

**Supplementary Table 1. Antibiotic sequencing analysis**

| Source | Chrm loc | Reference (MOY1355) | Mutation | Gene name | Location | Synonymous? |
|---|---|---|---|---|---|---|
| MOY1355 Cam$^R$ | IV 1456574 | T | TAAAAAC | SPG3 | I41* | No |
| | XII 6057 | A | T | | | |
| | Mito 16081 | CAATTA | C | AI1 | 756 | No |
| | Mito 61920 | A | C | 21S rRNA | A2769C | |
| MOY1355 Ery$^R$ | IV 1485641 | G | T | SPS1 | N466K | No |
| | VI 107508 | G | T | HSP12 | A85S | No |
| | VII 218009 | C | A | NUT1 | | Yes |
| | VII 441650 | G | A | SCW11 | 420 | Yes |
| | XV 1003903 | G | GT | | | |
| | Mito 59966 | A | G | 21S rRNA | A1958G | |

Supplementary table 2. Synthesized *RML2* genes

| Name | Sequence | Notes |
|---|---|---|
| *RML2* chimera with *S. cerevisiae* and *S. pombe* | GCATGGTCTCTAAAGATGTTGGTCTTGGGTTCTTTGAGATCTGCTTTGTCTTGTTCTTCTACCG<br>CCTCTTTGATTTCTAAGAGAAATCCATGTTACCCCTACGGTATTTTGTGTAGAACTTTGTCCCA<br>ATCCGTTAAGTTGTGGCAAGAAAACACTTCCAAGGATGACTCCTCTTTGAATATTACTCCAAG<br>GCTGTTGAAGATCATCCCAAACGATACTGATATCGTCACCTTGGAAAAGCAAGACGAATTGAT<br>CAAGCGTAGACGTAAGTTGTCTAAAGAAGTCACCCAAATGAAGAGATTGAAGCCAGTTTCTC<br>CAGGTTTGAGATGGTATAGATCTCCAATCTATCCCTACTTGTACAAAGGTAGACCAGTTAGAG<br>CTTTGACCGTTGTTAGAAAGAAACATGGTGGTAGAAACAACTCCGGTAAGATTACTGTTAGA<br>CATCAAGGTGGTGGTCATAGAAACAGAACTAGATTGATCGATTTCAACAGATGGGAAGGTGG<br>TGCTCAAACTGTTCAAAGAATTGAATATGACCCAGGCAGGTCATCTCATATTGCTTTGTTGAA<br>ACATAACACCACCGGTGAGTTGTCCTACATTATTGCTTGTGATGGTTTAAGACCAGGTGATGT<br>TGTTGAATCTTTCAGAAGAGGTATTCCACAGACCTTGTTGAACGAAATGGGTGGTAAAGTTG<br>ATCCAGCTATCTTGTCTGTTAAGACTACTCAAAGAGGTAACTGCTTGCCAATTTCCATGATTCC<br>AATTGGTACTATCATCCACAACGTTGGTATTACACCAGTTGGTCCAGGTAAATTCTGTAGATCT<br>GCTGGTACTTATGCTAGAGTTTTGGCTAAATTGCCAGAAAAGAAGAAGGCCATCGTTAGATT<br>ACAATCCGGTGAACATAGATACGTTTCTTTGGAAGCTGTTGCTACCATTGGTGTTGTCTCTAAT<br>ATCGATCACCAGAACAGATCTTTGGGTAAAGCTGGTAGATCTAGATGGTTGGGTATTAGACC<br>AACTGTTAGAGGTGTTGCTATGAACAAATGTGATCATCCTCATGGTGGTGGAAGAGGTAAAT<br>CTATTGGTAACAAACCTTCTCAATCTCCATGGGGTGTTTTGGCAAAAGGTGGTTACAAAACTA<br>GAAGGGGTAAGAACGTTAACAAGTTGTTGGTTAGAGATAGGCCTAGGGGTAAAGAAAAGAG<br>ATAAAAGAGACCGCAT | Highlighted sequence indicates region diverted from *S. cerevisiae RML2* gene.<br><br>Highlighted blue sections are added restriction sites for cloning. |

| | | |
|---|---|---|
| *RML2* chimera with *S. cerevisiae* and mitochondrial node | GCATGGTCTCTAAAGATGTTGGTCTTGGGTTCTTTGAGATCTGCTTTGTCTTGTTCTTCTACCG<br>CCTCTTTGATTTCTAAGAGAAATCCATGTTACCCCTACGGTATTTTGTGTAGAACTTTGTCCCA<br>ATCCGTTAAGTTGTGGCAAGAAAACACTTCCAAGGATGACTCCTCTTTGAATATTACTCCAAG<br>GCTGTTGAAGATCATCCCAAACGATACTGATATCGTCACCTTGGAAAAGCAAGACGAATTGAT<br>CAAGCGTAGACGTAAGTTGTCTAAAGAAGTCACCCAAATGAAGAGATTGAAGCCAGTTTCTC<br>CAGGTTTGAGATGGTATAGATCTCCAATCTATCCCTACTTGTACAAAGGTAGACCAGTTAGAG<br>CTTTGACCGTTGTTAGAAAGAAACATGGTGGTAGAAACAACTCCGGTAAGATTACTGTTAGA<br>CATCAAGGTGGTGGTCATAGAAACAGAACTAGATTGATCGATTTCAACAGATGGGAAGGTGG<br>TGCTCAAACTGTTCAAAGAATTGAATATGACCCAGGCAGGTCATCTCATATTGCTTTGTTGAA<br>ACATAACACCACCGGTGAGTTGTCCTACATTATTGCTTGTGATGGTTTAAGACCAGGTGATGT<br>TGTTGAATCTTTCAGAAGAGGTATTCCACAGACCTTGTTGAACGAAATGGGTGGTAAAGTTG<br>ATCCAGCTATCTTGTCTGTTAAGACTACTCAAAGAGGTAACTGCTTGCCAATTTCCATGATTCC<br>AATTGGTACTATCATCCACAACGTTGGTATTACACCAGTTGGTCCAGGTAAATTCTGTAGATCT<br>GCTGGTACTTATGCTAGAGTTTTGGCTAAATTGCCAGAAAAGAAGAAGGCCATCGTTAGATT<br>ACAATCCGGTGAACATAGATACGTTTCTTTGGAAGCTGTTGCTACCATTGGTGTTGTCTCTAAT<br>ATCGATCACCAGAACAGATCTTTGGGTAAAGCTGGTAGATCTAGATGGTTGGGTATTAGACC<br>AACTGTTAGAGGTGTTGCTATGAACAAATGTGATCATCCTCATGGTGGTGGAAGAGGTAAAA<br>CTTCAGGTGGTAGACCATCAGTTACTCCATGGGGTTGGCCTACAAAAGGTAAAAAGACTAGA<br>TCTAAGAGAGCCTCCGATAAGTTCATAGTCCAAAGAAGAAAGAAGAGGAAGAAGTAAAAGA<br>GACCGCAT | Highlighted yellow sequence indicates region diverted from *S. cerevisiae RML2* gene.<br><br>Highlighted blue sections are added restriction sites for cloning. |
| Codon-optimized *RML2* from *S. cerevisiae* | GCATGGTCTCTAAAGATGTTGGTCTTGGGTTCTTTGAGATCTGCTTTGTCTTGTTCTTCTACCG<br>CCTCTTTGATTTCTAAGAGAAATCCATGTTACCCCTACGGTATTTTGTGTAGAACTTTGTCCCA<br>ATCCGTTAAGTTGTGGCAAGAAAACACTTCCAAGGATGACTCCTCTTTGAATATTACTCCAAG<br>GCTGTTGAAGATCATCCCAAACGATACTGATATCGTCACCTTGGAAAAGCAAGACGAATTGAT<br>CAAGCGTAGACGTAAGTTGTCTAAAGAAGTCACCCAAATGAAGAGATTGAAGCCAGTTTCTC<br>CAGGTTTGAGATGGTATAGATCTCCAATCTATCCCTACTTGTACAAAGGTAGACCAGTTAGAG<br>CTTTGACCGTTGTTAGAAAGAAACATGGTGGTAGAAACAACTCCGGTAAGATTACTGTTAGA<br>CATCAAGGTGGTGGTCATAGAAACAGAACTAGATTGATCGATTTCAACAGATGGGAAGGTGG<br>TGCTCAAACTGTTCAAAGAATTGAATATGACCCAGGCAGGTCATCTCATATTGCTTTGTTGAA<br>ACATAACACCACCGGTGAGTTGTCCTACATTATTGCTTGTGATGGTTTAAGACCAGGTGATGT<br>TGTTGAATCTTTCAGAAGAGGTATTCCACAGACCTTGTTGAACGAAATGGGTGGTAAAGTTG<br>ATCCAGCTATCTTGTCTGTTAAGACTACTCAAAGAGGTAACTGCTTGCCAATTTCCATGATTCC<br>AATTGGTACTATCATCCACAACGTTGGTATTACACCAGTTGGTCCAGGTAAATTCTGTAGATCT<br>GCTGGTACTTATGCTAGAGTTTTGGCTAAATTGCCAGAAAAGAAGAAGGCCATCGTTAGATT<br>ACAATCCGGTGAACATAGATACGTTTCTTTGGAAGCTGTTGCTACCATTGGTGTTGTCTCTAAT<br>ATCGATCACCAGAACAGATCTTTGGGTAAAGCTGGTAGATCTAGATGGTTGGGTATTAGACC<br>AACTGTTAGAGGTGTTGCTATGAACAAATGTGATCATCCTCATGGTGGTGGAAGAGGTAAAT<br>CTAAATCTAACAAGCTGTCTATGTCTCCATGGGGTCAATTGGCTAAAGGTTACAAAACTAGAA<br>GGGGTAAGAATCAGAACAGGATGAAGGTTAAGGACAGGCCTAGAGGTAAGGATGCTAGATT<br>GTAAAAGAGACCGCAT | Highlighted blue sections are added restriction sites for cloning. |

Supplementary Table 3. Oligonucleotides

| Name | Sequence | Purpose |
| --- | --- | --- |
| prBRM001_gRNA seq FP | AAACAACTGTGTGATCCTTGAG | Fw and Rv primers for gRNA sequencing |
| prBRM002_gRNA seq RP | CCTGAGGAATCTTTAATACATTTTC | |
| prBRM041_L2_seqF | GTATGGGAAACTGAAAGCGAG | Fw and Rv primers for *RML2* ORF sequencing |
| prBRM042_L2_seqR | TTCAATATTTATATTTACATGATTGTGGAC | |
| prBRM043_L2_F1_F | TTTGGTCTCCCGCACTTACTAATAAGCGATGCTGGTTTTAGA GCTAGAAATAGCAAGTTAAAATAAGGCTAGTCCG | Fw and Rv primers for *RML2* frag 1 gRNAs (grey homology to Addgene plasmid 198866) |
| prBRM044_L2_F1_R | TTTGGTCTCCGTCTTTTACTTTTGCGCAAGCCGGGAATCGA | |
| prBRM045_L2_F2_F | TTTGGTCTCGAGACAGACCAAGGTTTTAGAGCTAGAAATAG CAAGTTAAAATAAGGCTAGTCCG | Fw and Rv primers for *RML2* frag 2 gRNAs (grey homology to CRISPR plasmid) |
| prBRM046_L2_f2_R | TTTGGTCTCGGGATTGCGCAAGCCGGGAATCGA | |
| prBRM062_L2_FP2 | TCATTTCAAAGATCCGCTTG | Fw and Rv primers for colony PCR verification of *RML2* insertion |
| prBRM063_L2_RP2 | TGGTTGCTGATCAAGAAGAC | |
| MRPL13-amp5 | CTCTTAACTCTTGTTCCCTGGC | Fw and Rv primers for *MRPL13* truncation verification |
| MRPL13-amp3 | TACGCAACTGGAGAGGAAGG | |
| MRP20-KanMX4-D1 | ATTCGACGGAGTCGTGGGGCCTTACGAACGGGTGGCCCAGTA GCGTACGCTGCAGGTCGAC | Deletion of 225-263 from *MRP20*, *KanMX4* from pKL55, black has gene homology, green is stop codon and red has pKL55 homology for PCR |
| MRP20-KanMX4-D2 | TTTGCGTGTGGGTTAACGAAGGGGTGGAATGGGAAACCTTAT CGATGAATTCGAGCTCGT | |
| MRPL13-URA3-D1 | AAGTAAATATGATACTATTATGAAAGAAATTCAAAAGTTATAG GCAGCAGATCTGGCTTTTC | Deletion of 250-264 from *MRP20*, *URA3* from pFL34, black has gene homology, green is stop codon and red has pFL34 homology for PCR |
| MRPL13-URA3-D2 | TAAAATGTAATGATAGTAGTAATAATAGTATTAACATAAGCAG ATCTGAGCTTTTTCTTTCC | |
| MRP20-amp5 | GTGAGCTTGTCTACGTTATTGG | Fw and Rv primers for *MRP20* truncation verification |
| MRP20-amp3 | CGTAAGGCATGTTCTACAGCC | |
| prBRM095_gR9_FP | TTTGGTCTCCCGCATTTTTCTGTAAACTCTTTATGTTTTAGAGCT AGAAATAGCAAGTTA | Fw and Rv primers for *RRG9* 39 gRNAs |
| prBRM096_gR9_RP | TTTGGTCTCCGGATTGCGCAAGCCGGGAATCG | |
| prBRM103_R9_FP2 | TTCCCTTCGACTTCCTTCTG | Fw and Rv primers for *RRG9* sequencing |
| prBRM104_R9_RP2 | GCAAGATCAGATGCTGTCAG | |
| pBRM075_R9_HA_FP | ATGGTCATCCAAAAAGATTATTAAATTAGTCAATAAATCGAtTT TGTCgAATAAAGAGTT | Fw and rv primers for homology arms and integration of Rrg9 S39I SNP |

| | | |
|---|---|---|
| pBRM076_R9_HA_RP | CTTTCGTACCATCTCGTACTTTTTCTGTAAACTCTTTATTcGACAAAaTCGATTTATTGA | |
| pBRM079_180g_FP | TTTGGTCTCGCGCAATTTTTGAAAAGAGGACAACGTTTTAGAGCTAGAAATAGCAAGTTAAAATAAGGCTAGTCCG | *MRX1* 180 gRNAs |
| pBRM080_180g_RP | TTTGGTCTCGGGATTGCGCAAGCCGGGAATCGA | |
| prBRM101_Ms_FP | AGCAGATACAAAACTGGGGC | Fw and rv primers for *MRX1* sequencing |
| prBRM102_Ms_RP | TCTTCAAAACAAGTTCTCCG | |
| pBRM081_180H_FP | TCCAAATTAATCGCCAATGATGTTCAAAAATTTTTGAAAAtAGGACAACTGGAAAAAGCT | Fw and rv primers for homology arms and integration of Mrx1 R180I SNP |
| pBRM082_180H_RP | AGCTAGCCTAGCCAAAAAGACAGCTTTTTCCAGTTGTCCTaTTTTCAAAAATTTTTGAAC | |
| pBRM087_224g_FP | TTTGGTCTCCCGCATTCAATTGGCGAAAAAAATGGTTTTAGAGCTAGAAATAGCAAGTTAAAATAAGGCTAGTCCG | *MRX1* 224 gRNAs |
| pBRM088_224g_RP | TTTGGTCTCCGGATTGCGCAAGCCGGGAATCGA | |
| pBRM089_224H_FP | GTTGTGCAGTCCCAACAAAGTGCTGTAGATATCTTCAATTcGCGAAAAAAATGGGGAGTC | Fw and rv primers for homology arms and integration of Mrx1 W224S SNP |
| pBRM090_224H_RP | GATGGAATGTTGATCAATAGGGACTCCCCATTTTTTTCGCgAATTGAAGATATCTACAGC | |
